# Head-mounted microendoscopic calcium imaging in dorsal premotor cortex of behaving rhesus macaque

**DOI:** 10.1101/2020.04.10.996116

**Authors:** Anil Bollimunta, Samantha R. Santacruz, Ryan W. Eaton, Pei S. Xu, John H. Morrison, Karen A. Moxon, Jose M. Carmena, Jonathan J. Nassi

## Abstract

A major effort is now underway across the brain sciences to identify, characterize and manipulate mesoscale neural circuits in order to elucidate the mechanisms underlying sensory perception, cognition and behavior. Optical imaging technologies, in conjunction with genetically encoded sensors and actuators, serve as important tools toward these goals, allowing access to large-scale genetically defined neuronal populations. In particular, one-photon miniature microscopes, coupled with genetically encoded calcium indicators and microendoscopic gradient-refractive index (GRIN) lenses, enable unprecedented readout of neural circuit dynamics in cortical and deep subcortical brain regions during active behavior in rodents. This has already led to breakthrough discoveries across a wide array of rodent brain regions and behaviors. However, in order to study the neural circuit mechanisms underlying more complex and clinically relevant human behaviors and cognitive functions, it is crucial to translate this technology to non-human primates. Here, we describe the first successful application of this technology in the rhesus macaque. We identified a viral strategy for robust expression of GCaMP, optimized a surgical protocol for microendoscope GRIN lens insertion, and created a chronic cranial chamber and lens mounting system for imaging in gyral cortex. Using these methods, we demonstrate the ability to perform plug-and-play, head-mounted recordings of cellular-resolution calcium dynamics from over 100 genetically-targeted neurons simultaneously in dorsal premotor cortex while the macaque performs a naturalistic motor reach task with the head unrestrained and freely moving. The recorded population of neurons exhibited calcium dynamics selective to the direction of reach, which we show can be used to decode the animal’s trial-by-trial motor behavior. Recordings were stable over several months, allowing us to longitudinally track large populations of individual neurons and monitor their relationship to motor behavior over time. Finally, we demonstrate the ability to conduct simultaneous, multi-site imaging in bilateral dorsal premotor cortices, offering an opportunity to study distributed networks underlying complex behavior and cognition. Together, this work establishes head-mounted microendoscopic calcium imaging in macaque as a powerful new approach for studying the neural circuit mechanisms underlying complex and clinically relevant behaviors, and promises to greatly advance our understanding of human brain function, as well as its dysfunction in neurological disease.

**Highlights:** First demonstration of head-mounted microendoscopic calcium imaging in behaving macaque.

Surgical protocols developed for preparing the animal for calcium imaging, including virus injections to express GCaMP and chronic implantation of a GRIN lens to enable optical access to gyral cortex.

Proof of concept plug-and-play calcium imaging in behaving macaques with months long stable recording capability allowing populations of individual neurons to be tracked longitudinally.

Bilateral calcium imaging from dorsal premotor cortex exhibited dynamics selective to the animal’s direction of reach and allowed decoding of the animal’s motor behavior

## Introduction

Mesoscale neural circuits have increasingly become a key focus of studies investigating brain function in health and disease. The development of new methodologies to support these studies has rapidly accelerated over the last decade, with optical technologies, such as calcium imaging and optogenetics, quickly becoming mainstays of circuit neuroscience investigation (Jennings and Stuber, 2014; Hamel et al 2015; Deisseroth, 2015; Yang and Yuste, 2017). Calcium imaging, bolstered by the explosive development of molecular-genetic tools (Chen et al., 2013b; Luo et al., 2018), offers several important advantages over conventional electrophysiological approaches. Notably, calcium imaging enables the ability to record with high cellular sampling density from large populations of genetically- or anatomically-defined neurons and to longitudinally track individual neurons over time (Hamel et al., 2015; Yang and Yuste, 2017). These powerful capabilities, now routinely applied in mouse models, have already yielded tremendous new insights into the neural basis of essential brain functions.

Despite these important advances, a critical need still exists to deploy these new technologies beyond the mouse to larger animal model species of relevance to humans (O’Shea et al., 2017). Non-human primates (NHPs) are a particularly important model species in this regard, with brain structure and function, as well as complex cognitive and behavioral abilities, highly similar to that of humans (Capitanio and Emborg, 2008; Phillips et al., 2014; Roelfsema and Treue, 2014). Additionally, recent advances in genome editing are quickly making NHPs viable genetic models of human disease (Sato and Sasaki, 2018). Therefore, transfer of the latest optical technologies from rodents to behaving NHPs promises to play a key role in elucidating clinically-relevant neural activity correlates of healthy and aberrant human behavior. Successful application of calcium imaging in NHPs, however, has been slow to develop for a number of reasons. In particular, difficulties using conventional viruses to express genetically-encoded calcium indicators in the NHP brain (Sadakane et al., 2015a) and imaging artifacts caused by movements of the larger volume NHP brain (Trautmann et al., 2015; Choi et al., 2018) have proven most challenging. Additionally, NHPs have a more mature immune system as compared to rodents that requires sophisticated surgical strategies and neural implant hardware, and limitations exist on the overall number of animals that can be used for trial-and-error technology development (Phillips et al., 2014).

Recent efforts applying two-photon microscopy and GCaMP-based calcium imaging in NHPs have overcome some of these challenges by refining aspects of the surgical and viral preparation and imaging system (Sadakane et al., 2015a; Seidemann et al., 2016; Yamada et al., 2016; Santisakultarm et al., 2016; Ebina et al., 2018; Zeng et al., 2019; Li et al., 2017; Ju et al., 2018; Garg et al., 2019; Trautmann et al., 2019a). These efforts have so far relied on transparent glass or silicone windows implanted over the brain to enable imaging of cortex to depths less than approximately 500 μm (e.g. cortical layers 2 and 3). The substantial motion artifacts inherent in this approach can be partially mitigated via mechanical pressure applied to the brain via the imaging window. While conventional adeno-associated viruses (AAVs) encoding GCaMP have in some cases proven sufficient for in vivo imaging (Li et al, 2017; Ju et al., 2018; Trautmann et al., 2019a), novel viral strategies such as AAVs utilizing tetracycline-controlled transcriptional activation (e.g. Tet-Off) have proven more effective at driving adequate levels of GCaMP expression (Sadakane et al., 2015a; Garg et al., 2019). While two-photon calcium imaging through cranial windows has many merits, it also has several limitations including: (1) infection risk of using glass or silicone surface windows, which also have limited functional lifetimes (e.g. degradation of optical clarity over time) and require daily maintenance, (2) cumbersome daily alignment between the microscope and the brain in order to maintain a consistent field of view (FOV) and track neurons across sessions, (3) inability to image gyral cortex deeper than approximately 500 μm, sulcal cortical areas or subcortical structures residing even deeper in the brain, (4) inability to measure from multiple brain areas in true simultaneity with a single scanning laser beam, and (5) necessity for head restraint, thereby limiting natural behavior.

An alternative to two-photon imaging that is poised to overcome many of these challenges in NHPs is the miniature, integrated one-photon fluorescence microscope (Ghosh et al., 2011; Ziv et al., 2013; Chen et al., 2013a; Ziv and Ghosh, 2015; Resendez et al., 2016). The nVista miniscope (Inscopix, Inc.) weighs less than 2 grams and can be carried on the head of a mouse while it is moving and behaving naturally. Used in conjunction with custom-designed gradient-refractive index (GRIN) microendoscopic lenses of various lengths and diameters, the nVista miniscope allows neuroscientists to visualize the activity of genetically defined neurons in almost all regions of the mouse brain, including deep brain structures inaccessible to other large-scale recording technologies. The first proof of concept demonstration of miniscope calcium imaging in NHP was recently demonstrated in deep layers of primary motor cortex of behaving marmosets (Kondo et al., 2018). This study confirmed several important advantages of this technology as applied in the NHP, including microendoscopic (as opposed to surface window) imaging access to deep cortical layers beyond the current depth limits of two-photon imaging and the ability to perform imaging during more natural, free behavior (e.g. head-unrestrained arm reaches, unrestrained ladder climbing) as compared to two-photon imaging under head-fixed conditions. While this study marked an important first step toward miniscope calcium imaging applications in NHPs, the marmoset’s small brain and lissencephalic cortex allowed for a relatively straightforward and smooth translation of methods already established in rodent models. The degree to which these methods would generalize to other NHP species, particularly those with larger brains and a gyrencephalic cortex, such as the more commonly studied rhesus macaque, remained to be determined.

Here, we demonstrate for the first time microendoscopic calcium imaging with head-mounted miniscopes in behaving rhesus macaque. To do so, we developed a viral strategy to express GCaMP in superficial and deep cortical layers, surgical methods for GRIN lens microendoscope probe insertion, and implant hardware optimized for performing chronic calcium imaging in macaque gyral cortex. The resulting imaging implants proved to be low maintenance, with little risk of infection and high optical clarity maintained throughout the entire period of study (> 8 months). We performed plug-and-play, head-mounted recordings of cellular-resolution calcium dynamics from over 100 genetically-targeted neurons in dorsal premotor cortex (PMd) while the macaque performed a naturalistic motor reach task with the head unrestrained and freely moving. We observed neuronal calcium dynamics selective to the animal’s directions of reach and a diversity of such selectivity across the recorded population. This population activity was used to decode the animal’s reach direction on individual trials. These recordings were stable over several months, allowing us to repeatedly measure the selectivity properties of populations of individual neurons over time. Finally, we took advantage of the small footprint of the miniscope to mount two miniscopes and perform simultaneous, multi-site imaging in PMd bilaterally, allowing for the measurement of both contralateral and ipsilateral neuronal population calcium dynamics during individual right or left arm reaches. These proof of concept results demonstrating head-mounted microendoscopic calcium imaging in behaving macaque provide an important step forward toward applying this powerful technology in NHPs and set the stage for future applications aimed at studying the neural circuit mechanisms underlying complex behavior and higher cognitive function in a model species similar to humans.

## Results

### Surgical preparation for microendoscopic calcium imaging in macaque dorsal premotor cortex

In this study, we demonstrate for the first time microendoscopic calcium imaging with head-mounted miniscopes in behaving rhesus macaque (Figure 1). Preparing the animal for imaging requires several steps, including two survival surgeries, the first to deliver a virus to the brain region of interest for expression of a genetically-encoded calcium indicator (e.g. GCaMP; Chen et al., 2013b) and the second to chronically implant a microendoscopic GRIN lens for optical access to that same brain region (Supplementary Figure 1).

**Figure 1.**
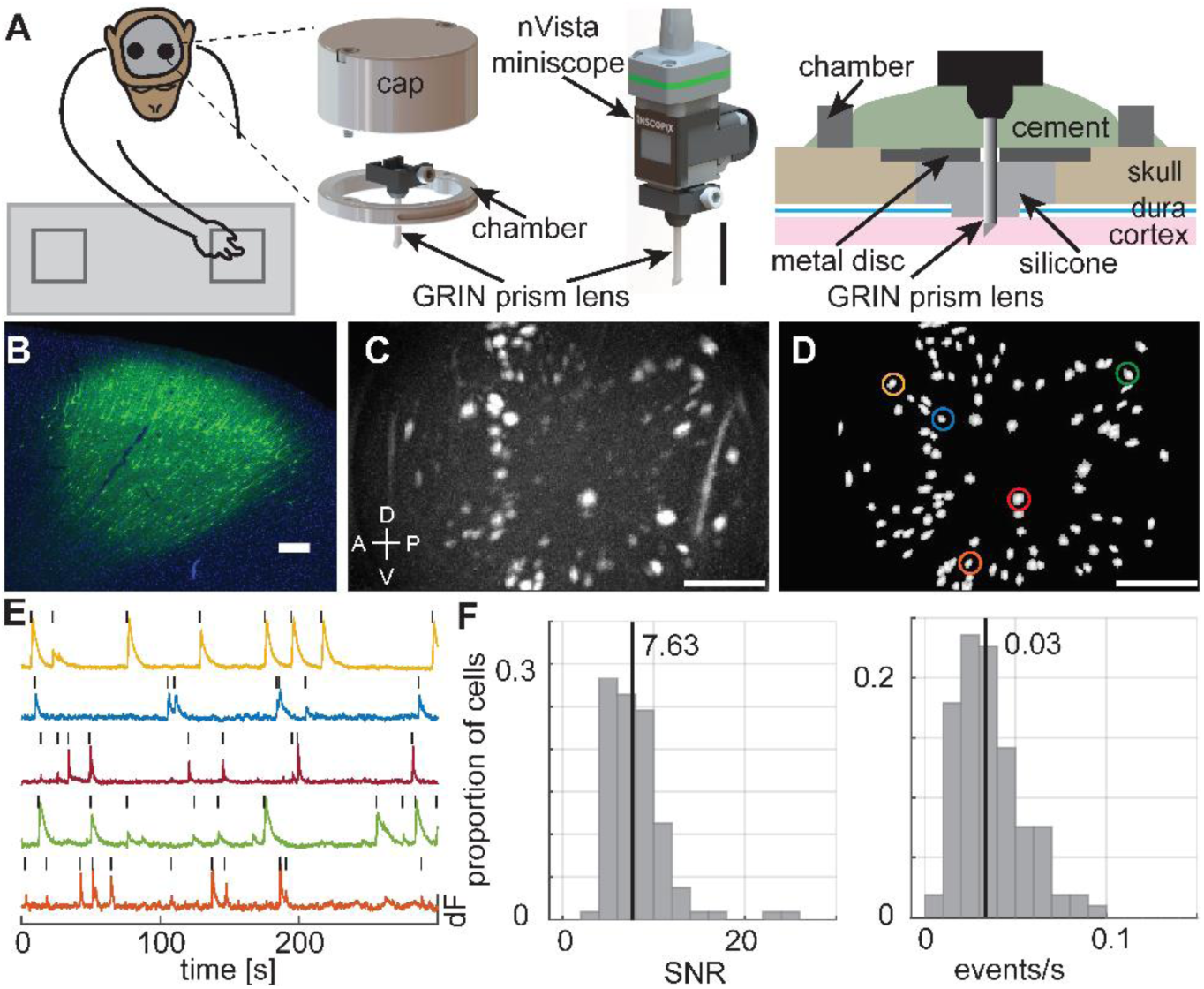
Cellular resolution imaging in macaque dorsal premotor cortex. **(A)** Left: Schematic of the macaque performing the reach to reward task with two nVista miniscopes mounted on the head to image from bilateral PMd. Middle: Zoomed in schematic of the implant hardware, including the GRIN prism lens integrated with the miniscope baseplate and the cranial chamber and cap, and the nVista miniscope docked on the baseplate for imaging. Scale bar equals 10 mm. Right: A schematic depicting how the chamber and lens probe were positioned and secured with respect to the cortex and skull. **(B)** Post-mortem native GCaMP expression (green) and DAPI-stained cell nuclei (blue) in the cortex 8 weeks following injections of the AAV Tet-Off virus system in a separate animal (animal 1). Scale bar equals 250 μm. **(C)** Max projection image of in vivo GCaMP fluorescence over the course of a single example session. The bright colored regions in the image indicate cells that exhibited active calcium dynamics during the recording. Scale bar equals 250 μm. Dorsal (D), Ventral (V), Anterior (A), Posterior (P) denote orientation in the premotor cortex. **(D)** Map of cells extracted using CNMFe from the same example session. Colored circles indicate example cell calcium activity traces in (E). Scale bar equals 250 μm. **(E)** Calcium activity (dF, peak normalized) traces of example cells highlighted in (D). The black tick marks above the traces indicate detected calcium events. **(F)** Distribution of median calcium event SNR (left) and median calcium event rate (right) for the entire population of cells recorded in the example session. The vertical lines indicate the median SNR (7.63) and event rate (0.03) values.

For calcium indicator expression in macaque cortex, we utilized two different AAV-based strategies tested first in a separate animal (Figure 1B, Supplementary Figure 2). One strategy used a conventional AAV (AAV1.CaMK2a.GCaMP6f) which has previously been shown to enable GCaMP-based two-photon calcium imaging in macaque cortex (Li et al., 2017). The second strategy used an AAV Tet-Off virus system consisting of two viruses mixed together (AAV5.Thy1s.tTA and AAV5.TRE3.GCaMP6f), which has been shown to express higher levels of GCaMP in shorter amounts of time as compared to conventional AAVs (Sadakane et al., 2015a; Garg et al., 2019). Furthermore, expression of GCaMP with this system is TET-dependent and therefore administration of Doxycycline (Dox) to the animal can temporarily suppress expression of GCaMP (Sadakane et al., 2015a; Kondo et al., 2018), which may be important for preventing overexpression for long-term chronic imaging. Both of these viral approaches resulted in robust cortical expression of GCaMP6f in the two animals we studied, with adequate spread across and within supragranular and infragranular layers of cortex (Supplementary Figures 2-3). The morphology of GCaMP-expressing cells indicated expression biased toward excitatory pyramidal neurons in the case of the CaMK2a virus and unbiased, pan-neuronal expression in the case of the Tet-Off virus system. Consistent with previous reports (Sadakane et al., 2015a; Kondo et al., 2018), the Tet-Off virus system resulted in higher levels of expression as compared to the conventional CaMK2a virus at the same time point post-injection (approximately 45% higher fluorescence intensity in post-mortem tissue; see Methods for details).

**Figure 2.**
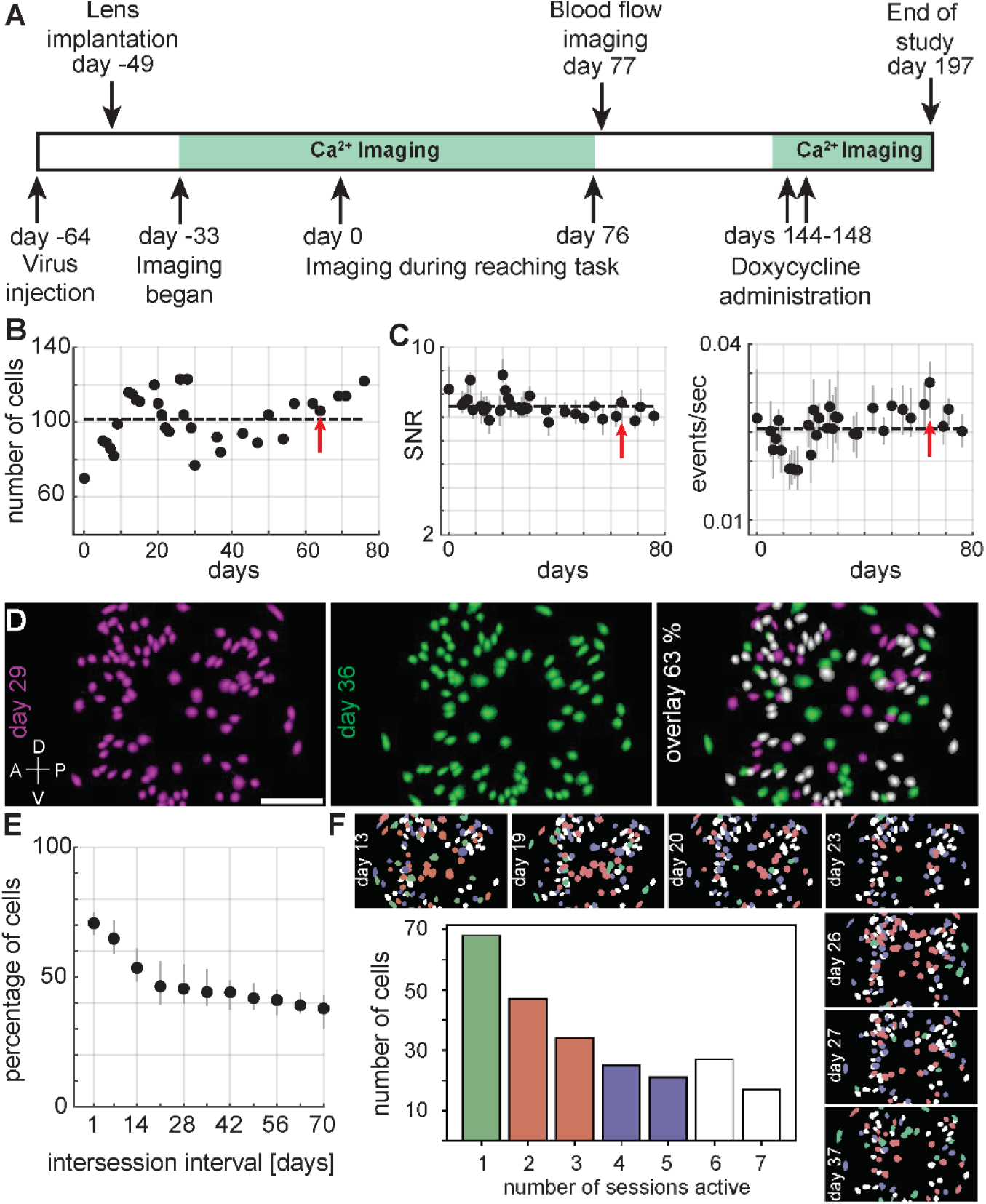
Calcium imaging stability and longitudinal tracking of neurons. **(A)** Schematic of the experimental timeline. **(B)** Number of cells that were imaged for each session across approximately 2.5 months (days 0 to 76 on experimental timeline in (A)). The dashed line indicates the mean value. The red arrow indicates the example session. **(C)** Calcium event SNR (left) and rates (right) (median and IQR) for each session across 76 days. The dashed line indicates the mean value. The red arrow indicates the example session. **(D)** Left: CNMFe-extracted cell map from an imaging session conducted on day 29 (magenta); Middle: cell map from day 36 (green); Right: Overlay of cell maps from the two sessions spaced 7 days apart. 63% of the cells from day 29, colored in white, were present and active on both days. Dorsal (D), Ventral (V), Anterior (A), Posterior (P) denote orientation in the premotor cortex. Scale bar equals 250 μm. **(E)** Percentage of cells (median, IQR) in common between two sessions as a function of the intersession interval (days). **(F)** Longitudinal tracking of cells through multiple sessions. Top and right side: cell maps from seven different sessions spanning approximately three weeks with individual cells color-coded according to the number of sessions in which they were present and active (colors as indicated in bar plot). Center: Percentage of cells as a function of the number of sessions (non-consecutive) found to be present and active.

We targeted the forelimb region of PMd bilaterally for imaging, relying on structural MRI images along with a standard atlas to determine stereotaxic coordinates prior to surgery, with minor adjustments to the final coordinates based on perioperative visualization of sulcal/gyral landmarks (see Supplementary Figure 3 for post-mortem confirmation). In the left hemisphere PMd we co-injected both viruses (mixed 1:1) comprising the Tet-Off virus system (titers: 2.18E+11 GC/ml for AAV5.mThy1PSs.tTAad and 2.08E+11 GC/ml for AAV5.TRE3.GCaMP6f) and in the right hemisphere PMd we injected AAV1.CaMK2a.GCaMP6f (titer: 2.3E+13 GC/ml) (see Methods for further details). In order to enable precise alignment of the GRIN lenses to the virus injection sites in the subsequent surgery (∼2 weeks later), we implemented two critical procedures: (1) photo documentation of the virus injection sites relative to the local blood vessel pattern and placement of fiducial marks on the skull, aligned with the virus injection sites, along with measurement of the distances between fiducial marks and the sites of injection, and (2) complete sealing of the durotomies using a combination of artificial dura and silicone elastomer, which prevented substantial neovascularization and pia/arachnoid cell proliferation during the time between surgeries (Supplementary Figure 1). Together, these two procedures allowed us to reference the fiducial marks and the local blood vessel patterns for precise alignment of virus injection and lens implantation sites. Prior to lens insertion, we made a linear incision in the cortex to create a path for the lens probe (Gulati et al., 2017; Kondo et al., 2018). We then inserted a GRIN prism lens (1 mm diameter, 9 mm length Proview Integrated Prism Lens; Inscopix, Inc.) approximately 2 mm below the surface of the cortex and at an angle perpendicular to the cortical surface, with the prism attached to the end relaying a side-view imaging plane (∼750 × 900 μm FOV) parallel to the axis of the probe and perpendicular to cortical layers (Supplementary Figure 3) (Gulati et al., 2017; Kondo et al., 2018). We chronically secured the top of the lens along with the integrated miniscope baseplate (which acts as a docking station for the miniscope) to the skull by embedding it in a combination of acrylic and cement (Figure 1A). We surrounded both lens implants with custom cranial chambers, each with a removable cap, that were embedded together with skull screws in a single acrylic headcap. Given that the craniotomies were completely sealed with cement/acrylic, the resulting implant had a low risk of infection and required little to no maintenance. Once the above two surgical steps were complete, the animal was ready for calcium imaging.

**Figure 3.**
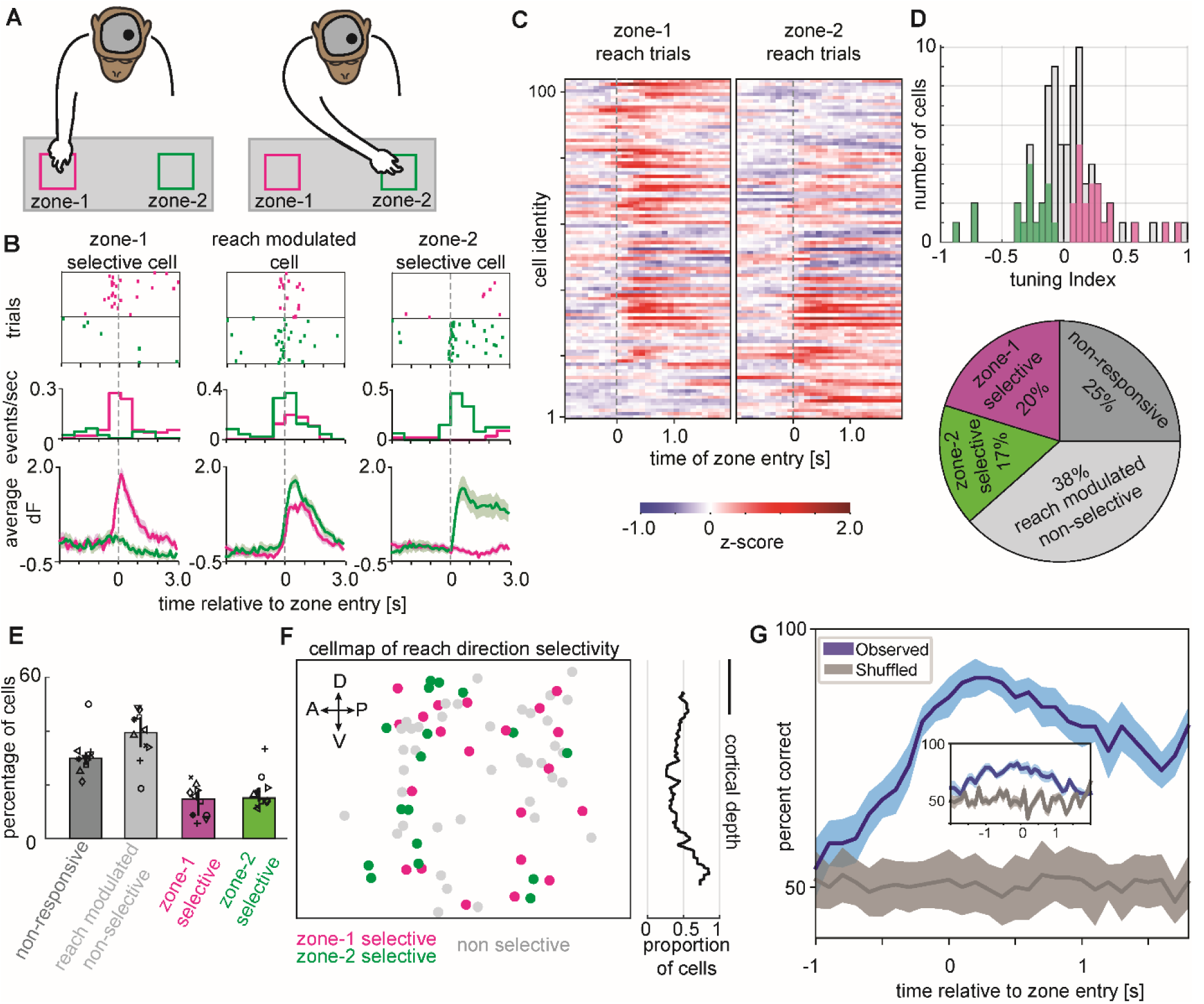
Direction selective calcium dynamics and decoding of motor reach behavior. **(A)** Schematic of the macaque performing the reach to reward task with an nVista miniscope mounted on the head to image from left hemisphere PMd. In these sessions, the macaque reached with the right arm (contralateral to the imaged hemisphere) to one of two zones, either zone 1 (magenta) or zone 2 (green). **(B)** Three example cells from the left hemisphere PMd exhibiting zone 1 selectivity (left), zone 2 selectivity (right) or nonselective modulation to either reach location (middle) in a single example session. Top: Rasters of calcium event times across multiple trials aligned to the time of reach entry (dashed vertical line) into zone 1 (magenta) or zone 2 (green). Middle: Peri-stimulus time histogram (PSTH) of calcium events as a function of time relative to reach entry into zone 1 (magenta) or zone 2 (green). Bottom: Calcium trace activity (mean, SEM) as a function of reach entry into zone 1 (magenta) or zone 2 (green). **(C)** Heatmap depicting z-scored trial-averaged calcium trace activity for each cell in the population (rows) on either zone 1 (left) or zone 2 (right) reach trials and aligned to the time of zone entry (dashed vertical line). The cells have been sorted top to bottom based on their selectivity (tuning index) to zone 1 or zone 2 reaches respectively. **(D)** Top: Distribution of reach direction selectivity (tuning index) for the entire population of cells recorded in the example session. Magenta and green colored bars indicate cells that had significant (p < 0.05; see Methods) reach direction selectivity to zone 1 (positive tuning indices) or zone 2 (negative tuning indices) respectively. Bottom: Pie chart depicting the percentage of cells in the example session that were classified as zone 1 selective (magenta), zone 2 selective (green), reach modulated but nonselective (light grey) or nonresponsive (dark grey). **(E)** Barplot depicting the median number of cells across sessions that are non-responsive (dark grey), task modulated but not reach direction selective (light grey), selective to zone-1 (magenta) and selective to zone-2 (green). Error bars are IQR. **(F)** Left: Cell map depicting the spatial distribution of reach direction selectivity. Cells selective for zone 1 (magenta), zone 2 (green) or reach modulated but nonselective (light grey) are indicated. Dorsal (D), Ventral (V), Anterior (A), Posterior (P) orientations in the left hemisphere PMd are indicated with arrows. Right: Proportion of reach modulated cells that were significantly selective to either reach direction as a function of cortical depth. Scale bar equals 200 μm. **(G)** Observed accuracy of decoding the animal’s reach direction on individual trials (mean, SEM) utilizing a model trained with calcium trace activity in 400 ms time bins (and 100 ms steps) around the time of reach entry into zones 1 and 2 (blue). Chance level decoding accuracy estimated by shuffling the reach direction across trials (grey). Inset: Decoding accuracy utilizing a model trained with calcium events instead of calcium traces.

### Plug-and-play, cellular resolution calcium imaging in alert macaque dorsal premotor cortex

Approximately two weeks following the lens implantation surgery, we initiated bilateral calcium imaging sessions in the alert macaque. A major advantage of the head-mounted miniscope and the implant design approach described here is that it enables plug-and-play imaging sessions (Supplementary Video 1). The animal, sitting comfortably in a standard primate chair, was temporarily restrained for miniscope docking, in this case with soft cushions placed against the sides of the head. In three simple and quick steps the cranial chamber cap was taken off, a cover on the miniscope baseplate was removed, and the miniscope (nVista 3.0; Inscopix, Inc.) was docked to the baseplate and secured in place by tightening a single set screw. The entire procedure was completed in less than 2 minutes per chamber. The animal’s head was then unrestrained and imaging commenced.

We observed GCaMP-expressing cells and corresponding cellular calcium dynamics in the very first imaging session conducted 2 weeks following lens implantation, and the number of cells ramped up to relatively stable values 4 weeks later when we started imaging while the animal performed a motor reach behavioral task (Figure 1C-F, Supplementary Video 2, Supplementary Figure 4). During these imaging sessions, the animal’s head was completely unrestrained and free to move naturally, and the animal was allowed frequent chewing of the food reward (Supplementary Video 2). Despite these significant jaw, head and body movements, the imaging FOV was very stable, requiring only a standard, rigid translation motion correction algorithm (Inscopix Data Processing Software, IDPS; Inscopix, Inc.) to precisely register frames across the entire recording (Supplementary Video 3). During a typical session, across all recorded video frames, the applied correction had a median of 0.75 μm (Interquartile range (IQR) = 0.5-1.1) which was an order of magnitude smaller than the imaged size of an average cell body. A small correlation between head movement and FOV translation was completely mitigated following motion correction (Pearson’s correlation: ρ = 0.11, p<0.001, before motion correction; ρ = 0.01, p=0.2, after motion correction; see Methods for further details).

**Figure 4.**
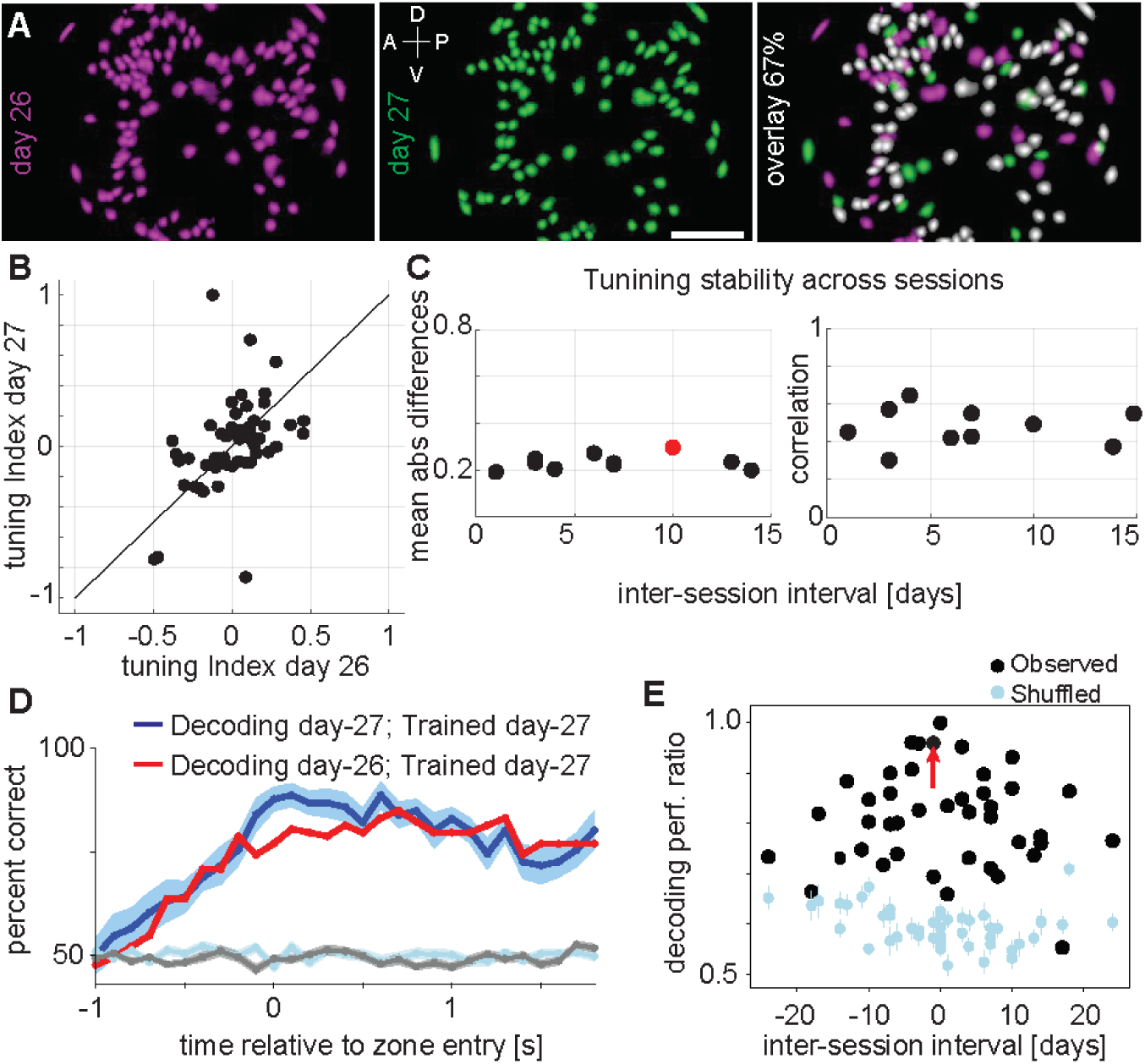
Longitudinally tracking the relationship between neurons and motor reach behavior. **(A)** Left: CNMFe-extracted cell map from an imaging session conducted on day 26 (magenta); Middle: cell map from day 27 (green); Right: Overlay of cell maps from the two sessions spaced 1 day apart. 67% of the cells from day 26, colored in white, were present and active on both days. Scale bar equals 250 μm. Dorsal (D), Ventral (V), Anterior (A), Posterior (P) denote orientation in the premotor cortex. **(B)** Reach direction selectivity (tuning index) on day 26 versus day 27 for cells exhibiting significant (p < 0.05; see Methods) reach direction selectivity to zone 1 (positive tuning indices) or zone 2 (negative tuning indices) and imaged longitudinally across both sessions. **(C)** Stability of reach direction selectivity across sessions. Left: Mean absolute differences in tuning index for cells imaged longitudinally across two sessions as a function of the inter-session interval (days). Sessions exhibiting significant differences indicated (red; Wilcoxon signed-rank test, p < 0.05). Right: correlation between tuning indices for cells imaged longitudinally across two sessions as a function of the inter-session interval (days). All correlations were found to be significant (p < 0.05). **(D)** Observed accuracy of decoding the animal’s reach direction on individual trials (mean, SEM) utilizing a model trained and tested on the same session (blue; day 27) or trained and tested on sessions 1 day apart (red; days 26 and 27). Chance level (across session) decoding accuracy estimated by shuffling the reach direction across trials (cyan) or by shuffling cell identity labels (grey). **(E)** Peak observed (black) and shuffled (cyan) decoding accuracy (mean, SEM) as a function of the inter-session interval (days). All values are expressed as a ratio normalized by the peak observed decoding accuracy for within-session training and testing. The red arrow points to the session pair in panel (D). See Supplementary Table 1 for decoding performance values for all sessions pairs.

We used constrained nonnegative matrix factorization (CNMFe), a common algorithm for cell identification from one-photon microendoscopic calcium imaging data, to identify individual cells and extract their cellular calcium dynamics (see Methods for more details) (Zhou et al., 2018). In the example left hemisphere recording shown (129 days post virus injection and 113 days post lens implantation), we identified 106 individual cells (Figure 1C, D). The calcium events had sufficient signal-to-noise (SNR) to be detected by a standard event detection algorithm (IDPS; Inscopix, Inc.), and had the typical and expected shape with a fast rise and exponential decay (rise median 0.20 s [IQR 0.15-0.22]; decay median 0.35 s [IQR 0.30-0.50]) (Figure 1E). Both SNR (median 7.6 [IQR 5.7-9.4]) and event rates (median 0.03 events/s [IQR 0.02-0.04]) measured across the population for this example session (Figure 1F) were similar to those typically observed in calcium imaging studies in rodent models (Chen et al., 2013b; Resendez et al., 2016). We observed significantly fewer imageable cells in the right hemisphere PMd, likely due to non-optimal placement of the lens relative to the zone of GCaMP-expressing cells (Supplementary Figures 3-4). Nevertheless, the detected calcium events measured from these cells had similar SNR (median 7.5 [IQR 6.8-8.4]) and event rates (median 0.03 events/s [IQR 0.02-0.05]) to that observed from cells in the left hemisphere.

### Chronic, longitudinal calcium imaging and tracking of populations of individual neurons over time

We performed a total of 66 imaging sessions over approximately 8 months, with 42 consecutive calcium imaging sessions over the first 4 months (Figure 2A, day 33 to day 76), followed by a pause in calcium imaging in order to conduct sedated blood flow imaging (Supplementary Video 4; see Methods for more details). Resumption of calcium imaging began one month later after the systemically injected fluorescein dye, which interfered with our ability to detect GCaMP fluorescence signals, completely dissipated. We continued calcium imaging during month 6 in order to test the effects of Dox administration on measured calcium dynamics. During the 32 imaging sessions from day 0 to day 76 (Figure 2A), where the animal performed the task satisfactorily, the overall imaging quality and measured calcium dynamics were found to be stable (Figure 2B-C, Supplementary Figure 4D-E). The number of cells identified in the left hemisphere PMd fluctuated from session to session (minimum = 70, maximum = 123), but was relatively stable with a median of 104 [IQR 91-113] imaged cells per session (Figure 2B). The SNR (median 7.4 [IQR 7.2-7.6]) and rates (median 0.026 events/s [IQR 0.024-0.028]) of detected calcium events were also relatively stable across sessions (Figure 2C). Similar stability was observed for cells imaged in the right hemisphere PMd (Supplementary Figure 4D-E).

A comparison of calcium event decay kinetics from left and right hemisphere cell populations across sessions indicated significantly slower decay in the left hemisphere compared to the right (left: median 0.67 s [IQR 0.56-0.76], right: median 0.23 s [IQR 0.22-0.24]; Wilcoxon rank-sum test, p<0.001), which, given equivalent SNR across the two hemispheres (left: median 7.35 [IQR 7.15-7.61], right: median 7.56 [IQR 7.09-7.78]; Wilcoxon rank-sum test, p=0.54), suggested that GCaMP expression levels were higher in the left hemisphere cell population than the right hemisphere cell population (see Discussion). This is also consistent with the post-mortem histology results (Supplementary Figure 2) which indicate that the Tet-Off virus system (injected into the left hemisphere of the imaged animal) resulted in higher levels of expression compared to the CaMK2a virus (injected into the right hemisphere of the imaged animal) at the same time point post-injection. Given that overexpression of GCaMP can eventually result in epileptiform activity and cellular toxicity (none of which were observed in this study) (Chen et al., 2013b; Resendez et al., 2016; Steinmetz et al., 2017), it is important for long-term chronic imaging experiments to be able to keep expression levels within a reasonable range. Therefore, we additionally tested the effects of Dox administration (5 days SID, 15 mg/kg) on measured calcium dynamics (Supplementary Figure 5, see Methods for more details). As expected, administration of Dox resulted in a significant reduction of GCaMP expression levels, as can be seen in the max projection images and quantified by full-FOV average fluorescence. The number of identified cells was reduced from 75 to 0 just 3 days following cessation of Dox (8 days after initiation of Dox). Dox administration also led to faster decay kinetics of measured calcium events, providing additional evidence for reduced GCaMP levels within cells. The full-FOV average fluorescence, number of identified cells and decay kinetics all returned to pre-Dox levels approximately 40 days following cessation of Dox (Supplementary Figure 5). In the final imaging session conducted 8 months after the first session, imaging quality was maintained and calcium dynamics and events rates remained comparable to those measured in the first 4 months of recordings. These results indicate that the Tet-Off viral strategy for GCaMP expression allows active maintenance of indicator expression within acceptable levels over months of chronic imaging study.

**Figure 5.**
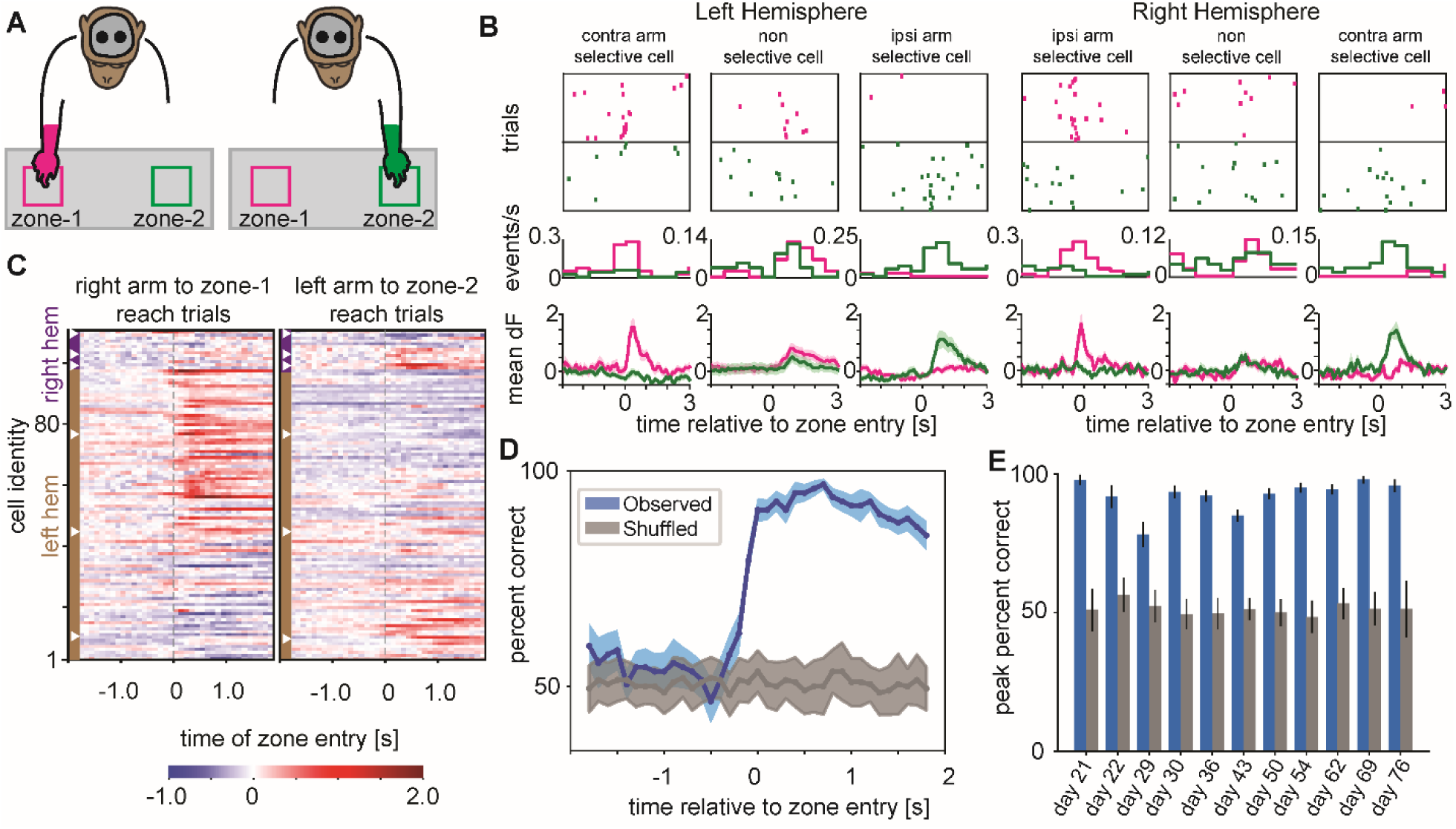
Multisite calcium imaging in bilateral dorsal premotor cortices with multiple head-mounted miniscopes. **(A)** Schematic of the macaque performing the reach to reward task with two nVista miniscopes mounted on the head to image bilaterally from left and right hemisphere PMd. In these sessions, the macaque reached with either the right arm to zone 1 (magenta) or the left arm to zone 2 (green). **(B)** Three example cells each from the left and right hemisphere PMd exhibiting right arm reach selectivity (left), left arm reach selectivity (right) or nonselective modulation to either arm reach (middle) in a single example session. Top: Rasters of calcium event times across multiple trials aligned to the time of reach entry (dashed vertical line) using right arm into zone 1 (magenta) or left arm into zone 2 (green). Middle: Peri-stimulus time histogram (PSTH) of calcium events as a function of time relative to reach entry using right arm into zone 1 (magenta) or left arm into zone 2 (green). Bottom: Calcium trace activity (mean, SEM) as a function of reach entry using right arm into zone 1 (magenta) or left arm into zone 2 (green). **(C)** Heatmap depicting z-scored trial-averaged calcium trace activity for each cell in the population (rows) on either right arm, zone 1 (left) or left arm, zone 2 (right) reach trials and aligned to the time of zone entry (dashed vertical line). The cells have first been grouped based on the hemisphere in which they reside (right hemisphere toward top [purple margin shading], left hemisphere toward bottom [brown margin shading]) and then within that group they have been sorted top to bottom based on their selectivity (tuning index) to right arm, zone 1 or left arm, zone 2 reaches respectively. The white arrowheads indicate the cells shown in (B). **(D)** Observed accuracy of decoding the animal’s reach arm on individual trials (mean, SEM) utilizing a model trained with calcium trace activity from bilateral PMd in 400 ms time bins (and 100 ms steps) around the time of right and left arm reach entry into zones 1 and 2 respectively (blue). Chance level decoding accuracy estimated by shuffling the reach arm across trials (grey). **(E)** Peak observed (blue) and shuffled (grey) decoding accuracy (mean, SEM) across sessions utilizing a model trained with calcium trace activity from bilateral PMd.

Given the overall stability of imaging quality and calcium dynamics across sessions, we next assessed how well we could track the same individual neurons across sessions, which is a major advantage of optical imaging techniques compared to electrophysiology (Ziv et al., 2013; Sheintuch et al., 2017). We demonstrated these capabilities for imaging in behaving macaque here, by applying a modified version of an established longitudinal cell registration algorithm (IDPS; Inscopix, Inc.) (see Methods for more details). For any two imaging sessions (7 day separation in Figure 2D), the cell maps extracted by CNMFe were first spatially registered by rigidly translating one map to the other in order to maximize the cross-correlation between them (Figure 2D). Once the cell maps for the two imaging sessions were registered, we calculated the spatial correlation between each cell and each of the rest of the cells in the population to detect candidate same-cell pairs across imaging sessions. Any cell pairs with a spatial correlation above a threshold value were considered putative same-cell pairs (colored white in Figure 2D). For the example session pair shown, we were able to identify 63% of cells (61 out of 97) imaged on day 29 as the same cells imaged on day 36 (see Supplementary Figure 6A for additional examples). Applying this approach to all possible imaging session pairs resulted in a median of approximately 70% same-cell pairs with minimum intersession intervals of 1-4 days, which dropped and plateaued to around 40% same-cell pairs with maximum intersession intervals of 67-73 days (Figure 2E). We further asked whether we could track the same populations of neurons through multiple, consecutive imaging sessions, rather than simply across any two session pairs. To track neurons across multiple sessions, we focused on a subset of 7 imaging sessions spanning approximately 3 weeks duration and applied a custom longitudinal registration algorithm (Figure 2F; see Methods for more details). We were able to track the same neurons across multiple imaging sessions, finding 68 cells active on only 1 of the 7 imaging sessions and 17 cells active in all 7 of 7 imaging sessions. As expected, the number of same-cell pairs identified across sessions, both for pairwise and continuous registration approaches, was dependent on the chosen spatial correlation criterion/threshold (Supplementary Figure 6B). Together, these results validate the ability to image the calcium dynamics of large populations of neurons in behaving macaque, over many months, with sufficient stability in imaging quality to longitudinally track individual neurons over time.

### Decoding of natural motor reach behavior from ensemble calcium dynamics

We next sought to directly relate population calcium related activity to the animal’s behavior. The animal was trained to perform a natural motor reach task in which the location of a food reward was randomly alternated across trials between two distinct locations (zone 1 and 2) and once the reward was placed in one of those locations the animal could reach, pick up and consume the food reward (Figure 3A). The animal made reaches with either the right or left arm in alternating blocks of trials. Based on previous electrophysiological studies in macaque PMd, we expected neural activity to be selective for the animal’s direction of reach, with particularly strong modulation of activity during reaches of the arm contralateral to the recorded hemisphere (Riehle and Requin, 1989; Kurata, 1993; Cisek and Kalaska, 2005). Our results confirmed these expectations, with clear evidence for cellular calcium dynamics selective for reach direction in both the left and right hemispheres (Figure 3B-D, Supplementary Video 5, Supplementary Figure 7). In the example left hemisphere recording shown (day 27, see Figure 2A), where the animal was reaching with the right arm, several cells exhibited increased calcium activity and a higher probability of calcium events during both zone 1 and zone 2 reaches, while other cells exhibited increased activity for zone 1 reaches compared to zone 2 reaches, or vice versa. Three example cells from this session clearly demonstrate these distinct selectivity profiles (Figure 3B), both in terms of reach-modulated calcium event rates and calcium trace dynamics.

We observed substantial diversity in reach-related modulation of calcium trace activity across the entire population of imaged cells in the example session (Figure 3C), with a majority of cells (75%, 78/104 cells) exhibiting significant modulation of activity associated with one or both reach directions (see Methods). We calculated a tuning index to capture the degree to which each cell was selective for a particular reach direction across the entire population (Figure 3D). We then used that tuning index to classify cells as either zone 1 selective, zone 2 selective or non-selective, finding that 35 of 78 cells were selective, 20% for zone 1 and 17% for zone 2. We applied this approach to individual sessions and found similar percentages of reach-modulated cells (median 67.8 [IQR 70.1-72.6]), and selectivity to zone 1 (median 14.7 [IQR 8.6-17.9]) and zone 2 (median 15.0 [IQR 13.7-18.9]), across sessions (Figure 3E). Using the same tuning index classification, we then mapped the distribution back on to the imaged FOV in the brain to assess whether selectivity for reach direction is spatially organized in PMd (Figure 3F). We did not observe any obvious spatial organization in these selectivity maps. To test if cells with similar selectivity tend to reside closer together than expected by chance, we calculated a cluster metric, which is the frequency with which the closest neighboring cell is similarly selective (see Methods). A cluster metric value that is significantly greater than 0.5 would suggest clustering of similarly selective cells. For the example session shown in Figure 3F the cluster metric was 0.57 and not significantly different from 0.5 (p = 0.16; bootstrap test). The GRIN prism lens used in the present study allows for simultaneous imaging of cells spread across multiple cortical layers. To investigate if the reach direction selectivity is more prevalent in specific cortical layers we estimated the proportion of direction selective cells as a function of cortical depth (Figure 3F, right). For the example session shown here the prevalence of selectivity was found to be fairly uniform across the depth of the cortex within the FOV.

Given the presence of reach location selective calcium activity in the recorded population, we investigated whether we could use the cellular calcium dynamics from the ensemble to decode the animal’s reach location on a trial-by-trial basis. Partial Least Squares-Discriminant Analysis (PLS-DA) with leave-one-out cross validation was used to decode reach direction from the continuous calcium trace activity of all identified cells within a 400 ms sliding window stepped at 100 ms (see Methods for more details). Decoding accuracy was well above chance and peaked around the time of zone entry (Figure 3G; see Supplementary Figure 7G for similar results from the right hemisphere). Decoding accuracy was stable across sessions (Supplementary Figure 8). Despite the sparseness of calcium events relative to continuous calcium traces, decoding accuracy based on calcium events was also well above chance (Figure 3G inset; see Methods for details on decoding algorithm used). As expected, decoding accuracy was dependent on the number of cells used to train the decoder, but remained above chance levels with as few as just 1 cell included for training (Supplementary Figure 8).

### Tracking the relationship between ensemble calcium dynamics and behavior over time

Having established that neurons in both left and right hemisphere PMd exhibited calcium dynamics selective for reach direction (Figure 3, Supplementary Figure 7) and having validated our ability to longitudinally track populations of neurons across sessions (Figure 2, Supplementary Figure 6), we were interested to determine the degree to which the direction selectivity of individual neurons changes or remains stable over time. We focused this analysis on a subset of left hemisphere imaging sessions for which the animal completed adequate numbers of reach trials, spanning 5 different sessions across 2 weeks duration. For the example session pair shown (Figure 4A, spaced 1 day apart), we spatially registered their cell maps and identified 82 putative same-cell pairs (123 total cells imaged on day 26 and 104 total cells imaged on day 27) (Figure 4A). We then compared the tuning indices for each of these same-cell pairs across the two imaging sessions, observing significant correlation (0.45, p < 0.001) and no significant change in the tuning of the population (Wilcoxon signed-rank test, p = 0.98) (Figure 4B). As inter-session interval increased, correlation between tuning indices across session pairs remained high and changes in tuning indices across session pairs remained low, indicating stable direction selectivity in PMd cells over greater than 2 weeks (Figure 4C).

Given the relatively stable population tuning dynamics measured for longitudinally tracked cells across sessions, we predicted that a decoder trained with calcium trace activity from a given session would perform reasonably well when tested on data from a separate session. This was confirmed for the same example session pair (spaced 1 day apart), where decoding accuracy was well above chance level and peaked around the time of zone entry, both for within-session testing and testing across the 1 day inter-session interval (Figure 4D). As inter-session interval increased, peak decoding accuracy remained above chance levels in most cases (Figure 4E, Supplementary Table 1), indicating that the encoding of reach direction in PMd is relatively stable over days to weeks. Together, these results provide proof of concept for how longitudinally tracked neurons can be applied toward advancing our understanding of the dynamic relationship between neural activity and behavior.

### Multi-site bilateral calcium imaging in dorsal premotor cortices

The results presented so far come from the left or right hemisphere PMd separately, while the animal reached with the arm contralateral to the recorded hemisphere. In recording sessions from day 20 to day 76 we additionally imaged simultaneously from both hemispheres (Figure 5). Due to the small size of the miniscope there was more than enough space on the head to mount two miniscopes, with ample space remaining for several more as may be desired in future studies. Despite imaging significantly fewer cells in the right hemisphere PMd as compared to the left hemisphere PMd (Figure 1, Supplementary Figure 4), these recordings provided an opportunity to establish proof of concept for simultaneous multi-site imaging. In order to take advantage of bilateral imaging capabilities, we trained the animal to perform a similar motor reach task to that already described, except the animal was trained to use the right arm to reach to zone 1 and the left arm to reach to zone 2 (Figure 5A). This task design, combined with simultaneous recordings from both the left and right hemisphere PMd, allowed us to investigate how arm reaches are bilaterally encoded on a trial-by-trial basis.

We found clear evidence for cellular calcium dynamics selective for ipsilateral or contralateral reach in both the left and right hemispheres (Figure 5B-C, Supplementary Video 6). In the example bilateral recording session shown in Figure 5 (day 29), several cells exhibited increased calcium activity and a higher probability of calcium events during both ipsilateral and contralateral reaches, while other cells exhibited increased activity for ipsilateral reaches compared to contralateral reaches, or vice versa. Three example cells from each hemisphere clearly demonstrate these distinct selectivity profiles (Figure 5B), both in terms of reach-modulated calcium event rates and calcium trace dynamics. We observed substantial diversity in reach-related modulation of calcium trace activity across the entire population of bilaterally imaged cells (Figure 5C). While most cells exhibited significant modulation of activity associated with contralateral reaches, a minority of cells were sensitive to ipsilateral reaches.

Given the presence of selectivity for both ipsilateral and contralateral reaches among the populations of recorded neurons in both hemispheres, we investigated whether we could use the cellular calcium dynamics from the bilateral ensemble to decode the identity of the arm being used to reach on a trial-by-trial basis. As before, we used PLS-DA with leave-one-out cross validation to decode left or right arm reach from continuous calcium trace activity within a 400 ms sliding window stepped at 100 ms. In this example session, decoding accuracy was well above chance and peaked around the time of zone entry (Figure 5D). Peak decoding accuracy was stable across sessions (Figure 5E). Together, these results establish proof of concept simultaneous multi-site imaging and provide preliminary data applying these capabilities toward advancing our understanding of the bilateral encoding of motor reach behavior.

## Discussion

We demonstrated here the first successful microendoscopic calcium imaging with head-mounted miniscopes in behaving rhesus macaque. As part of this work, we developed an effective strategy and surgical protocol for virally-mediated expression of GCaMP, as well as placement of a microendoscopic GRIN prism lens co-aligned to a region of interest in gyral cortex (Supplementary Figures 1-3). Our surgical approach, including the placement of fiducial marks on the skull and the complete sealing of the durotomy and craniotomy to preserve the health of the underlying cortex, proved very helpful for co-alignment. The complete implant, including the GRIN lens, integrated miniscope baseplate and cranial chamber, enabled plug-and-play daily recordings (Figures 1-2; Supplementary Figure 4; Supplementary Videos 1-3) and chronic, long-term study of naturalistic reaching behavior (Figures 3-5; Supplementary Figures 5-8; Supplementary Videos 5-6). We were able to measure the calcium dynamics of large ensembles of neurons (over 100 neurons per day on average in left hemisphere PMd) with SNR and event rates consistent with expectations from previous one-photon microendoscopic calcium imaging studies in rodents (Chen et al., 2013b; Resendez et al., 2016). These high quality imaging results were obtained with miniscopes mounted directly to the animal’s head with the head unrestrained and free to move naturally. Importantly, despite significant head movements, including chewing of rewards regularly throughout the recordings, we observed a highly stable FOV with only minor offline correction required. This approach, therefore, enables recording of neural activity during more natural behavioral conditions (e.g. head unrestrained), as compared to other large-scale recording technologies, greatly expanding the range and complexity of behaviors that can be studied (Gilja et al., 2010; Foster et al., 2014).

Our surgical and imaging approach allowed us to measure the calcium activity of neuronal ensembles in PMd cortex while simultaneously monitoring the animal’s motor reach behavior (Figure 3). We found evidence for reach-related modulation and selectivity preferences from the calcium traces and the extracted calcium events of recorded neurons (Figure 3B-C, Supplementary Figure 7). Furthermore, both the calcium traces and events could be used to decode the animal’s trial-by-trial direction of reach with accuracy significantly above chance levels, with highest performance levels using traces (Figure 3F, Supplementary Figure 8). The lower decoding performance for calcium events is likely related to their relative temporal sparsity, consistent with previous results (Ziv et al., 2013), and therefore we primarily utilized calcium traces for our analyses relating neural activity to behavior. It was striking to learn that high decoding performance could be maintained even when randomly sub-sampling just a single neuron from the population (Supplementary Figure 8), providing strong evidence for the encoding of motor reach behavior among the imaged ensemble. Importantly, the within-session selectivity preference distributions and decoding performances we observed in PMd are consistent with previously-reported results obtained electrophysiologically (Riehle and Requin, 1989; Kurata, 1993; Cisek and Kalaska, 2005). The ability to relate the functional properties of neuronal populations to their precise location and spatial organization within the brain is an additional major advantage of the optical imaging approach used here. We were able to map the selectivity preferences of neurons back on to the imaged FOV and assess the laminar distribution, and found overall heterogeneous spatial organization of selectivity for direction of reach (Figure 3F), which is again consistent with results obtained electrophysiologically (Trautmann et al., 2019b).

Given the time investment associated with training NHPs on complex cognitive and behavioral tasks, studies in macaque often involve experimentation in each individual animal lasting months to years. It is therefore critical that recording technologies applied in NHP have the longevity to support such chronic, long-term studies. Here, we showed that the number of recorded cells and the SNR and event rates measured from those cells were relatively stable across several months of recordings (Figures 2, 4). Despite most of these sessions occurring within a two month time window, we observed similar quality results in the very final session prior to discontinuing the study, suggesting longevity of at least 8 months and likely much longer. It is critical in such studies that GCaMP expression is maintained at reasonable levels, as overexpression of GCaMP can lead to cellular toxicity and abnormal epileptiform activity (Chen et al., 2013b; Resendez et al., 2016; Steinmetz et al., 2017). We did not observe any evidence for such effects of overexpression in either hemisphere and, although the decay kinetics of measured calcium events were longer in the left hemisphere as compared to the right (likely indicating higher levels of GCaMP in Tet-Off virus-injected left hemisphere PMd), they were nevertheless relatively stable across the entire duration of the study.

The overall stability across sessions in imaging quality and cellular GCaMP signals allowed us to longitudinally track large populations of individual neurons over several weeks to months. We note that the number of putative ‘same-cell’ pairs obtained from the longitudinal registration algorithm described here is sensitive to the inter-neuronal spatial correlation criterion chosen (Figure 2, Supplementary Figure 6). It will be critical, therefore, to develop improved methods for objectively and quantitatively determining the criterion/threshold in future studies (Sheintuch et al., 2017). Importantly, we show that the longitudinal tracking of neurons allows for the investigation of the functional properties of a circuit over time and the dynamic relationship between those functional circuit properties and the behaviors they subserve (Figure 4). While these analyses were only meant to be proof of concept applications utilizing longitudinally registered cell sets, the observed stability in neuronal selectivity dynamics, which is consistent with results obtained electrophysiologically (Chestek et al., 2007), and the ability to decode reach behavior in sessions days to weeks apart from the training session, constitute exciting results worth future study. These types of investigations are particularly valuable for understanding the functional plasticity of circuits in the context of learning and memory paradigms.

It is worth highlighting here several advantages of one-photon microendoscopic calcium imaging as compared to conventional electrophysiology that were confirmed as part of this study. As opposed to electrophysiological approaches that are typically blind to the neuronal subtype identity of recorded neurons (but see Movshon and Newsome, 1996; Lima et al., 2009; Kaufman et al., 2010; Kaufman et al., 2013; Mitchell et al., 2007; Trainito et al., 2019), optical imaging approaches combined with genetically encoded fluorescent activity indicators such as GCaMP can target and record specifically from a variety of anatomically and genetically-defined neuronal subtypes (Luo et al., 2018), with increasing options for similarly doing so in NHP models (Tervo et al., 2016; Dimidschstein et al., 2016; Stauffer et al., 2016; Mehta et al., 2019; Weiss et al., 2020). Here, we used two different viral-based strategies to genetically target and image either pan-neuronal or excitatory neuron-biased populations as a first step toward imaging of more specific cell populations. Another limitation of electrophysiological approaches, is the relatively sparse spatial sampling density of today’s commercially available multi-electrode arrays, typically having a minimum inter-electrode spacing of approximately 400 μm (Leber et al., 2019; but see Jun et al., 2017; Trautmann et al., 2019a for use of 20 μm spacing NeuroPixels in rhesus macaque). In contrast, cellular-resolution optical imaging enables much higher sampling densities, as evidenced by our study here where we were able to simultaneously record from more than 100 cells per session on average in the left hemisphere PMd (Figures 1-2), which equates to a sampling density of approximately 150 cells per mm^2^. This high sampling density allows for critical investigation of microcircuit organization and function typically off limits to electrophysiological approaches. Finally, as already discussed, we confirmed that this imaging approach enables the longitudinal tracking of populations of individual neurons over several months (Figure 2), which is either not possible or extremely challenging with traditional electrophysiological recordings that rely on waveform shape and timing characteristics alone for any attempted certification of neuron identity (Chestek et al., 2007; Ganguly and Carmena, 2009; Fraser and Schwartz, 2012).

This study also confirmed several advantages of one-photon microendoscopic calcium imaging with head-mounted miniscopes in comparison to traditional two-photon microscopy, especially as applied to NHP models. Most two-photon calcium imaging studies to date have relied on transparent cranial windows to optically access the superficial layers of cortex (Sadakane et al., 2015b; Li et al., 2017; Trautmann et al., 2019a). In NHPs, these cranial window implants can be challenging to maintain and are associated with a high risk of infection, typically exhibiting a degradation in optical clarity (a “clouding” of the implant) over time due to the proliferation of pia and arachnoid cells and, in some cases, the re-growth of native dura (Arieli et al., 2002; Chen et al., 2002). In our study here we confirmed sufficient optical clarity and quality of calcium imaging out to approximately 8 months and discontinued the study for reasons independent of implant integrity. We did not observe evidence of any degradation in optical clarity over the duration of the study and expect that maintenance of proper levels of GCaMP expression is the more critical consideration for chronic, long term microendoscopic calcium imaging studies in NHP. Due to the fact that the skull is completely sealed with cement surrounding the GRIN lens, the implant is very low maintenance (requires no cleanings or sterile procedures) and has a low additional risk of infection beyond the standard risks associated with invasive survival surgeries. An additional challenge associated with two-photon microscopy techniques relates to the often cumbersome daily setup and alignment between the microscope objective and the intended imaging plane, which even when refined requires non-negligible time (Choi et al., 2018; Trautmann et al., 2019a). In the study here we show that the implant supports remarkably easy miniscope docking and recording (Supplementary Video 1) and, in combination with the miniscope’s electronic focusing mechanism, enables precise alignment to the intended imaging plane across repeated sessions (Figure 2, Supplementary Figure 6).

Additional advantages over two-photon microscopy result from the small, head-mounted design of the miniscopes and their seamless coupling to microendoscopic GRIN lenses implanted deep in the brain. These features allowed us to mount two miniscopes on the head and record simultaneously from bilateral PMd (Figure 5). Conventional two-photon microscopy with a single scanning laser beam can only image from one site at a time, though some new approaches to two-photon are pushing these limits (Terada et al., 2018; Yang et al., 2019). The small footprint of the miniscope, in comparison, enables several brain regions to be imaged in true simultaneity. Because the miniscopes are all head-mounted, imaging does not require head fixation as is required for two-photon imaging studies. This, in turn, enables head unrestrained behavior during imaging, which is an important step toward more natural, freely behaving paradigms. Nevertheless, due to the miniscope cable, such studies still require that the macaque be partially restrained in a primate chair as is common practice in most NHP labs. Development of wireless miniscopes, which would enable fully unrestrained and free behavior, should be feasible using the hardware architectures already developed for comparable bandwidths, such as wireless transmission of high channel count electrophysiological recordings in NHPs (Miranda et al., 2010; Gao et al., 2012). Finally, the easy plug-and-play coupling of the miniscopes to the implanted microendoscopic GRIN lens enables access to brain regions deeper than are typically accessible from the brain surface with two-photon microscopy. In this study, post-mortem histological assessments indicated that the distal end of the probe was likely situated near the border of layers 3 and 4, at a max depth of approximately 1.5 mm below the cortical surface. These depths are well beyond that typically attainable with two-photon imaging from the surface.

Limitations of the imaging approach described here include potential damage to brain tissue and local circuits resulting from the implantation of a microendoscopic GRIN lens with relatively large diameter. These lenses are available in diameters ranging from 0.5 to 1.0 mm, with larger diameters (e.g. 1.0 mm used in this study) offering a larger imaging FOV, but with higher risk of tissue displacement or damage. Two-photon imaging through surface windows offers a less invasive approach, requiring no penetration into the brain, yet as already discussed is limited to depths less than approximately 500 μm, leaving most of the macaque brain inaccessible. Standard electrodes used for electrophysiological recordings in macaque can reach deep into the brain and are typically much thinner (e.g. 35-300 μm diameter), though one would need to implant many such electrodes in close proximity to approach the same number and density of cells that can be imaged with a single lens. Furthermore, the size of the GRIN lens used here compares with that of standard deep-brain stimulation (DBS) electrodes that have been used successfully in human patients and in animals with similarly sized brains (Miocinovic et al., 2013; Johnson et al., 2013). It is important to note that, except for the tissue damage created by the cortical incision made just prior to lens insertion, the actual lens itself may simply displace the remaining tissue without necessarily causing additional damage. More studies are needed to characterize the injury to the brain during implantation and the foreign body response to the chronically implanted lens (Barretto et al., 2011; Kozai et al., 2015; Lee et al., 2016). Another limitation associated with the use of a microendoscopic GRIN lens, particularly when combined with a one-photon miniscope, is the lower spatial resolution as compared to that of a two-photon microscope. While two-photon imaging through surface windows can often resolve static and dynamic signals at the scale of fine anatomical structures such as dendritic spines (Sadakane et al., 2015b; Li et al., 2017), one-photon microendoscopic imaging is typically limited to dynamic signals associated with larger structures such as cell bodies (Ghosh et al., 2011; Resendez et al., 2016). Finally, both one- and two-photon imaging currently rely on calcium indicators with relatively slow kinetics as a measurement of neural activity (Chen et al., 2013b), with significantly poorer temporal resolution as compared to electrophysiological measurements. However, the development of genetically encoded fluorescent voltage indicators (Villette et al., 2019; Knöpfel and Song, 2019) promises to overcome this limitation in the near future.

In order to take full advantage of head-mounted microendoscopic calcium imaging in behaving macaques, future studies should focus on several key areas of development. First, it is important to further streamline the surgical workflow, to enable virus injections and lens implantations functionally targeted and precisely aligned to each other and to the brain regions of interest. Virus-coated GRIN lenses (e.g. ProView Express Probes; Inscopix, Inc.) could completely obviate the need for alignment between separate virus injection and lens implantation surgeries, combining all components of the imaging preparation into a single surgery and a single penetration into the brain. Second, new viral strategies and longer GRIN lenses will be important for targeting specific cell-types and deeper brain regions respectively. Given the small size of the miniscope, it is possible to target several (beyond just two as shown here) brain regions simultaneously to investigate multiple nodes of the networks underlying complex behavior. As studies increasingly seek to record from the NHP brain in more naturally behaving conditions, development of a miniscope capable of wireless transmission will be critical. Finally, combining miniscope calcium imaging with optogenetic stimulation (e.g. nVoke system, Inscopix, Inc.; Stamatakis et al., 2018) or electrophysiology would allow for important studies testing the causal relationship between the functional properties of a circuit and the relevant behavior. Together with these future developments, microendoscopic calcium imaging with head-mounted miniscopes in NHPs will enable important new insights into the neural circuit mechanisms underlying clinically-relevant human behavior and will help to advance our understanding of and ability to develop effective therapies for neurodegenerative and neuropsychiatric disorders.

## Supplementary Figures and Table

**Supplementary Figure 1.**
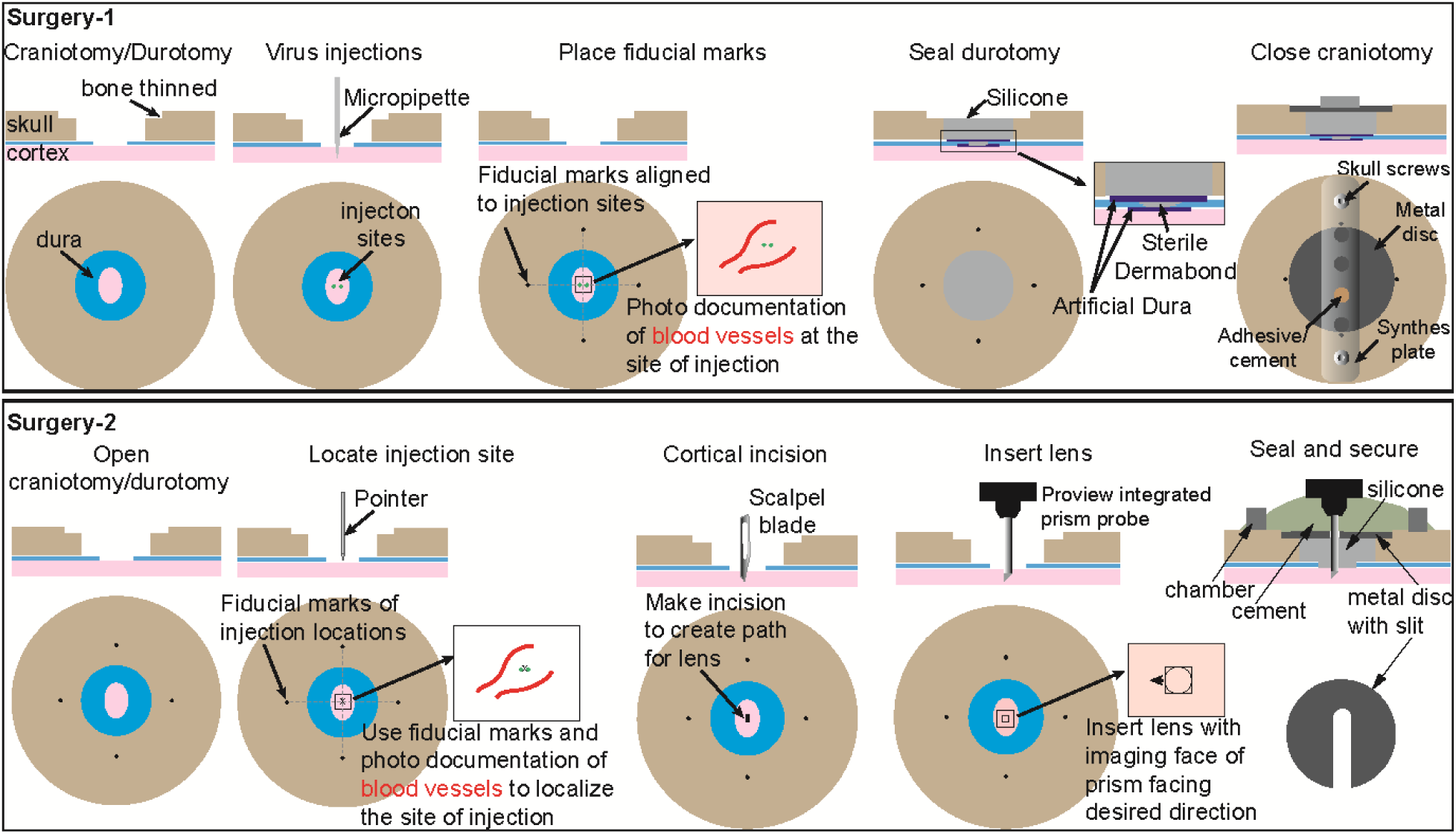
Schematic of surgical steps to prepare macaque for chronic imaging. Top: A schematic of surgical steps during the first surgery for injections of virus. Bottom: A schematic of surgical steps during the second surgery for lens implantation. See Methods for more details.

**Supplementary Figure 2.**
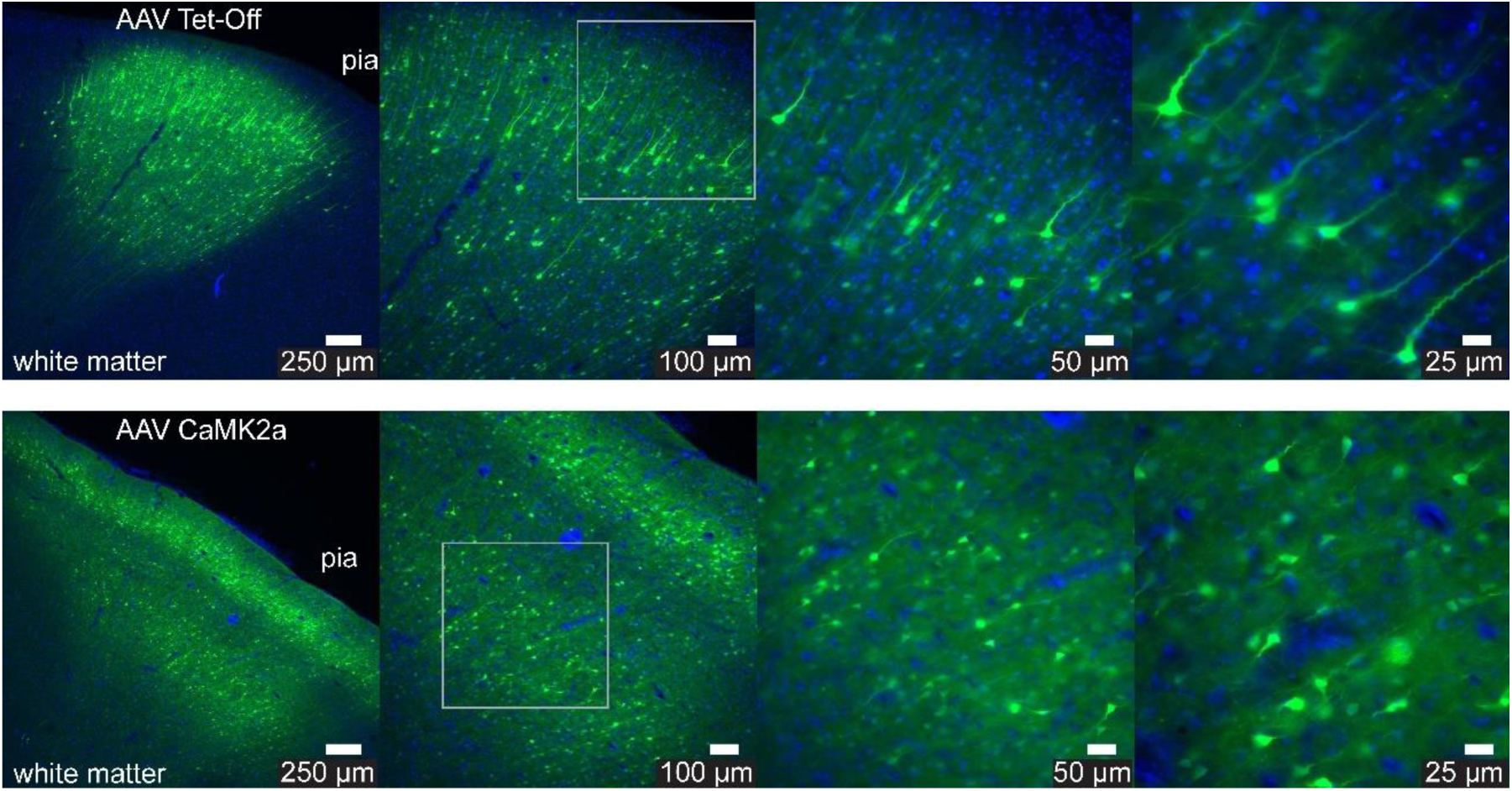
Post-mortem assessment of GCaMP expression in macaque 1. Top: Post-mortem native GCaMP expression (green) and DAPI-stained cell nuclei (blue) in the cortex 8 weeks following injections of the AAV Tet-Off virus system (top) or AAV1 CaMK2a virus (bottom) in animal 1. Images of increasing magnification are shown from left to right, with scale bars equal to 250, 100, 50 and 25 μm respectively. The higher magnification images to the right are from the superficial (top row) or deep (bottom row) layers indicated by the rectangle in the middle-left panel.

**Supplementary Figure 3.**
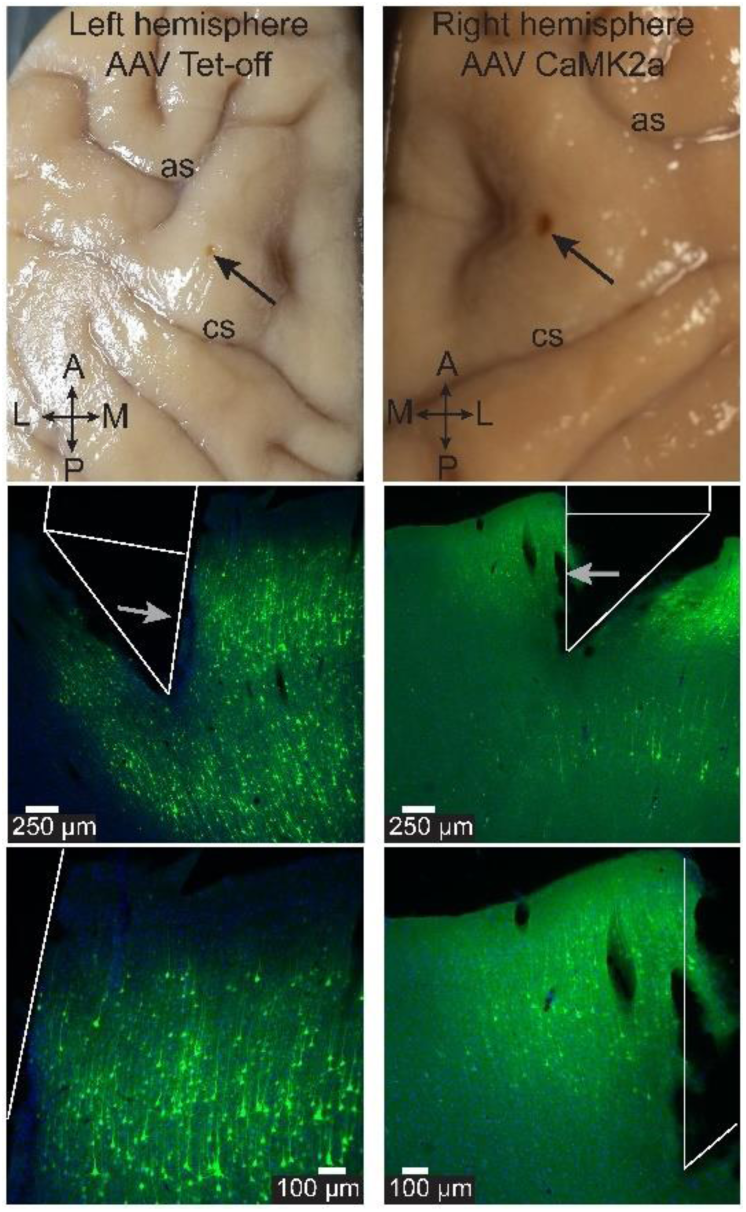
Post-mortem assessment of GCaMP expression and location of lens implants in macaque 2. Top: Post-mortem, ex vivo image of dorsal surface of brain and lens implant site for the left hemisphere (left; AAV Tet-Off) and right hemisphere (right; AAV CaMK2a) from animal 2. The arrow indicates the location of the lens implant. ‘as’ arcuate sulcus, ‘cs’ central sulcus. ‘A’ anterior, ‘P’ posterior, ‘M’ medial, ‘L’ lateral. Middle: Post-mortem native GCaMP expression (green) and DAPI-stained cell nuclei (blue) in PMd cortex 8.5 months following injections of the AAV Tet-Off virus system (left) or AAV CaMK2a (right) in animal 2. The estimated location of the prism lens during imaging is outlined in white. The arrow indicates the imaging surface of the prism and the direction of imaging. M-L axis as in the top panels. Scale bars equal 250 μm. Bottom: Same as above at higher magnification. Scale bars equal 100 μm.

**Supplementary Figure 4.**
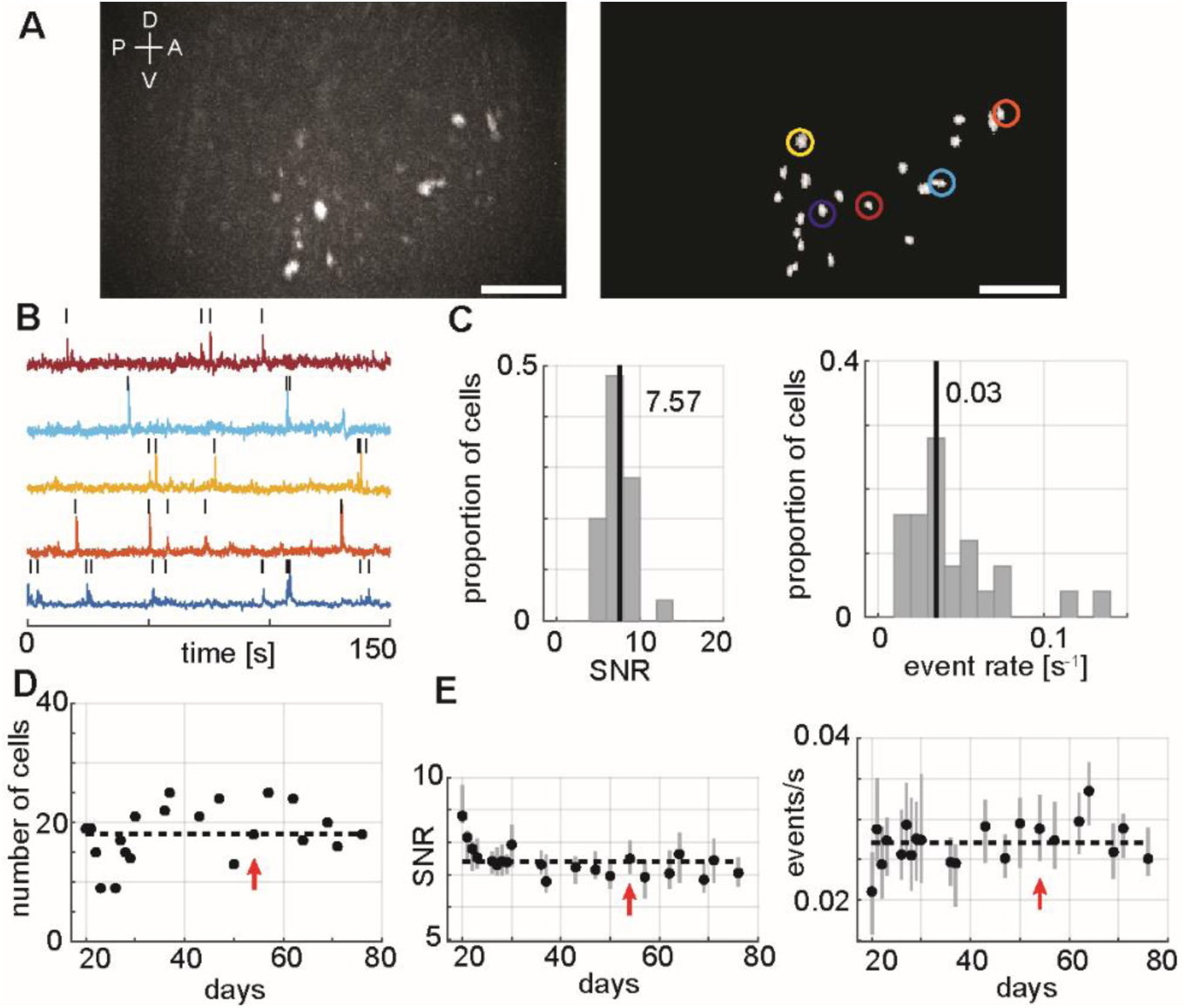
Cellular resolution imaging in macaque dorsal premotor cortex right hemisphere. **(A)** Left: Max projection image of in vivo GCaMP fluorescence in the right hemisphere PMd over the course of a single example session. The bright colored regions in the image indicate cells that exhibited active calcium dynamics during the recording. Dorsal (D), Ventral (V), Anterior (A), Posterior (P) denote orientation in the premotor cortex. Scale bar equals 250 μm. Right: Map of cells extracted using CNMFe from the same example session. Colored circles indicate example cell calcium activity traces in (B). Scale bar equals 250 μm. **(B)** Calcium activity (dF, peak normalized) traces of example cells highlighted in (A). The black tick marks above the traces indicate CNMFe extracted calcium events. **(C)** Distribution of median calcium event SNR (left) and median calcium event rate (right) for the entire population of cells recorded in the example session. The vertical lines indicate the median SNR (7.57) and event rate (0.03) values. **(D)** Number of cells that were imaged for each session across 60 days. The dashed line indicates the mean value. The red arrow indicates the example session. **(E)** Calcium event SNR (left) and rates (right) (median and IQR) for each session across 60 days. The dashed line indicates the mean value. The red arrow indicates the example session.

**Supplementary Figure 5.**
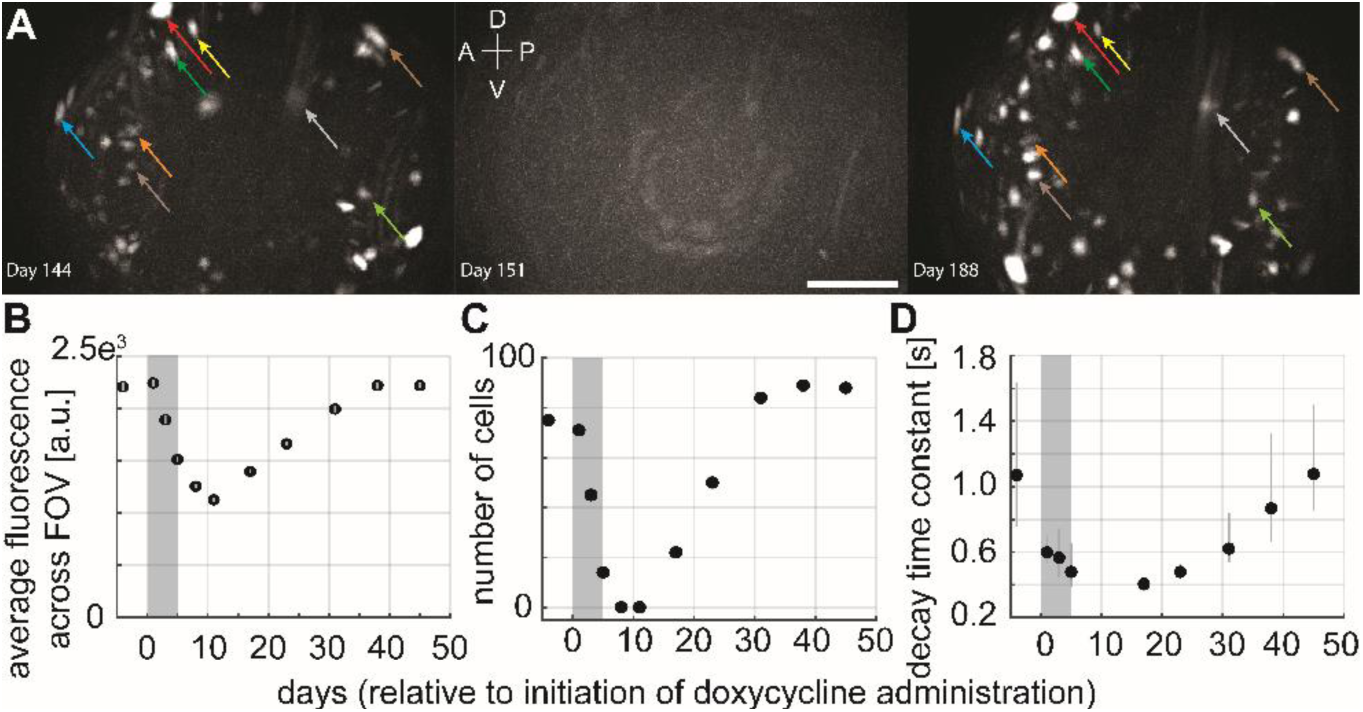
Control of Tet-Off virus mediated GCaMP expression with Doxycycline. **(A)** Max projection images of in vivo GCaMP fluorescence 1 day prior to initiation of Doxycycline (Dox) administration (left), 8 days following initiation (3 days following end) of Dox administration (middle) and 45 days following initiation (40 days following end) of Dox administration (right). Colored arrows indicate putative same-cells pairs identified 1 day prior to and 45 days following the initiation of Dox administration. Dorsal (D), Ventral (V), Anterior (A), Posterior (P) denote orientation in the premotor cortex. Scale bar equals 250 μm. **(B)** Average fluorescence across the FOV before, during and after Dox administration (mean, SEM). Grey shaded region indicates the time period of Dox administration. Fluorescence levels were reduced following Dox administration and returned to baseline approximately 35 days following its cessation. **(C)** Number of cells that were imaged for each session before, during and after Dox administration. Grey shaded region indicates the time period of Dox administration. The number of cells was reduced following Dox administration and returned to baseline approximately 30 days following its cessation. **(D)** Calcium event decay time constants (mean, 95% CI) before, during and after Dox administration. Grey shaded region indicates the time period of Dox administration. Decay time constants were reduced following Dox administration and returned to baseline approximately 40 days following its cessation.

**Supplementary Figure 6.**
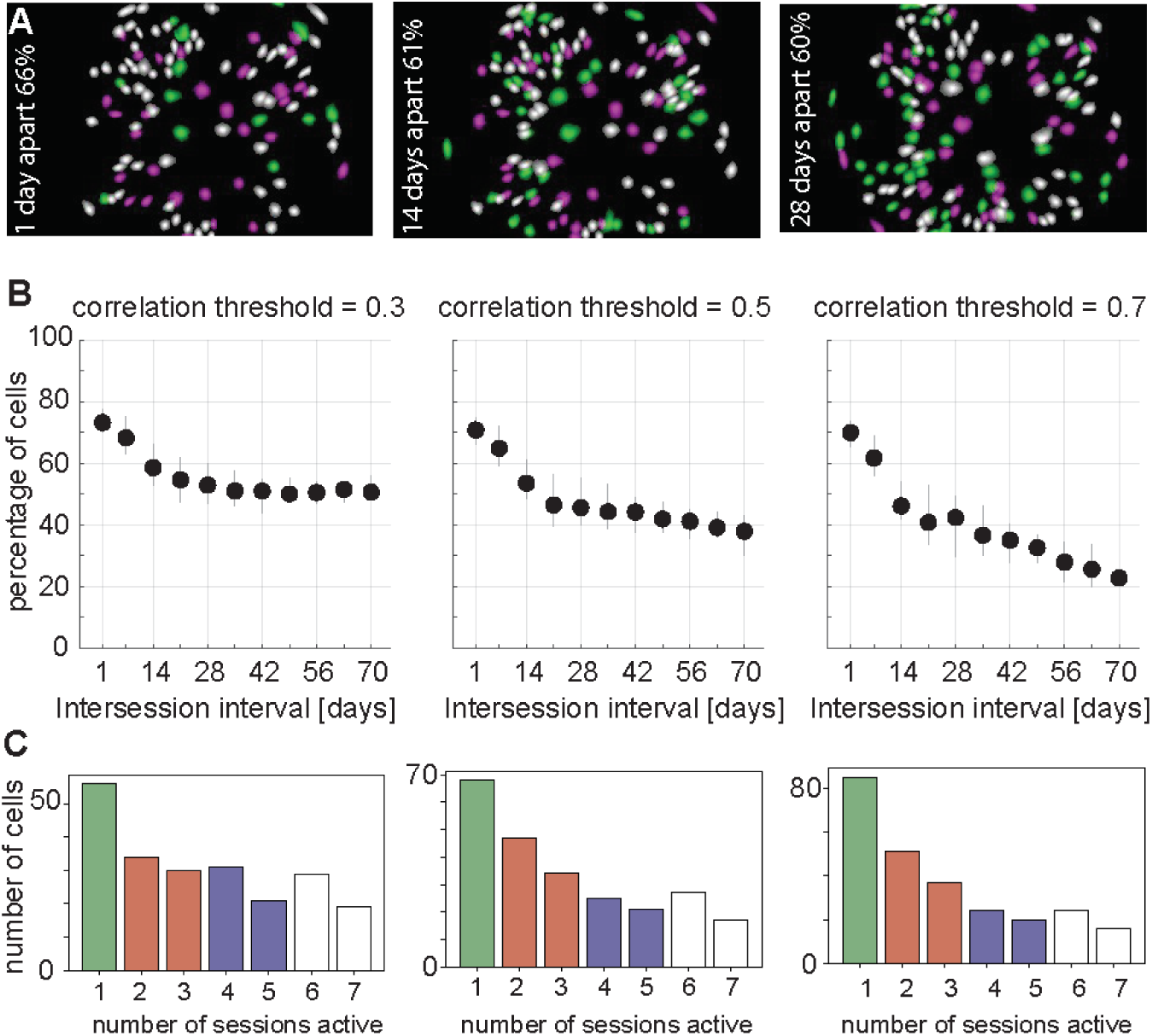
Dependencies of longitudinal tracking performance across sessions. **(A)** Overlays of CNMFe-extracted cell maps from two separate sessions (magenta and green cells) spaced 1 (left), 14 (middle) and 28 (right) days apart. The percentage of cells present and active in both sessions (white cells) is reported in each case. **(B)** Percentage of cells (median, IQR) in common between two sessions as a function of the intersession interval (days) for different spatial correlation thresholds used to determine putative ‘same-cell’ pairs (left = 0.3, middle = 0.5, right = 0.7). **(C)** Percentage of cells as a function of the number of sessions (non-consecutive) found to be present and active for different spatial correlation thresholds (left = 0.3, middle = 0.5, right = 0.7).

**Supplementary Figure 7.**
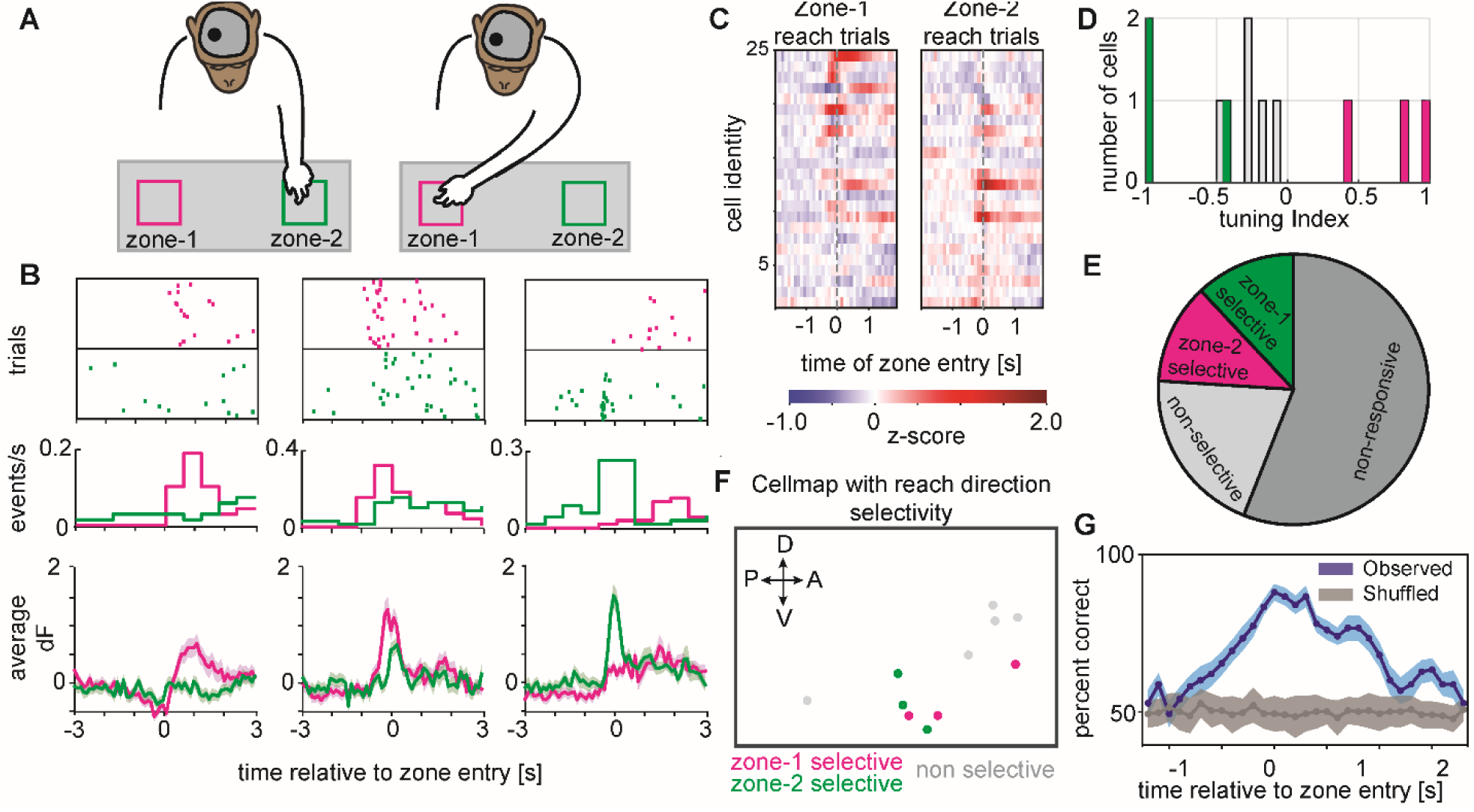
Direction selective calcium dynamics in right hemisphere PMd and decoding of motor reach behavior. **(A)** Schematic of the macaque performing the reach to reward task with an nVista miniscope mounted on the head to image from right hemisphere PMd. In these sessions, the macaque reached with the left arm (contralateral to the imaged hemisphere) to one of two zones, either zone 1 (magenta) or zone 2 (green). **(B)** Three example cells from the right hemisphere PMd exhibiting zone 1 selectivity (left), zone 2 selectivity (right) or nonselective modulation to either reach location (middle) in a single example session. Top: Rasters of calcium event times across multiple trials aligned to the time of reach entry (dashed vertical line) into zone 1 (magenta) or zone 2 (green). Middle: Peri-stimulus time histogram (PSTH) of calcium events as a function of time relative to reach entry into zone 1 (magenta) or zone 2 (green). Bottom: Calcium trace activity (mean, SEM) as a function of reach entry into zone 1 (magenta) or zone 2 (green). **(C)** Heatmap depicting z-scored trial-averaged calcium trace activity for each cell in the population (rows) on either zone 1 (left) or zone 2 (right) reach trials and aligned to the time of zone entry (dashed vertical line). The cells have been sorted top to bottom based on their selectivity (tuning index) to zone 1 or zone 2 reaches respectively. **(D)** Top: Distribution of reach direction selectivity (tuning index) for the entire population of cells recorded in the example session. Magenta and green colored bars indicate cells that had significant (p < 0.05; see Methods) reach direction selectivity to zone 1 (positive tuning indices) or zone 2 (negative tuning indices) respectively. **(E)** Pie chart depicting the percentage of cells in the example session that were classified as zone 1 selective (magenta), zone 2 selective (green), reach modulated but nonselective (light grey) or nonresponsive (dark grey). **(F)** Cell map depicting the spatial distribution of reach direction selectivity. Cells selective for zone 1 (magenta), zone 2 (green) or reach modulated but nonselective (light grey) are indicated. Dorsal (D), Ventral (V), Anterior (A), Posterior (P) denote orientation in the premotor cortex. **(G)** Observed accuracy of decoding the animal’s reach direction on individual trials (mean, SEM) utilizing a model trained with calcium trace activity in 400 ms time bins (and 100 ms steps) around the time of reach entry into zones 1 and 2 (blue). Chance level decoding accuracy estimated by shuffling the reach direction across trials (grey).

**Supplementary Figure 8.**
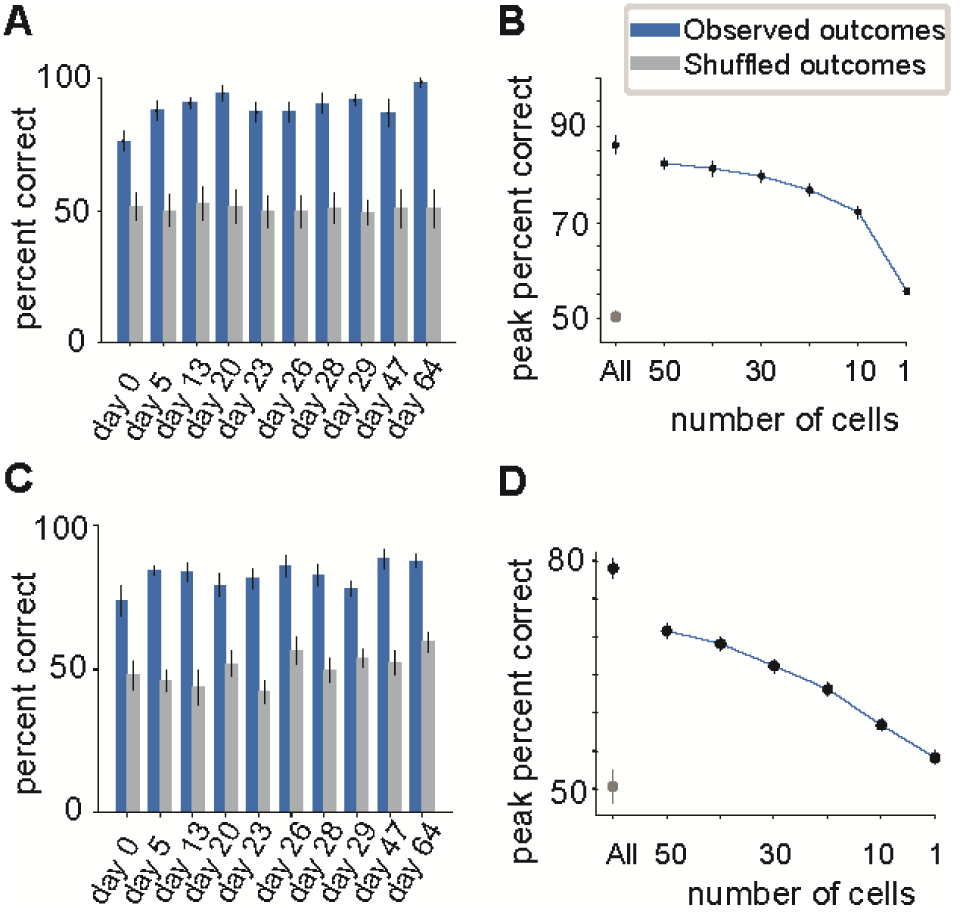
Decoding of motor reach behavior with calcium activity traces and events. **(A)** Peak observed (blue) and shuffled (grey) decoding accuracy (mean, SEM) across sessions utilizing a model trained with calcium trace activity. **(B)** Peak observed (blue) and shuffled (grey) decoding accuracy (mean, SEM) utilizing a model trained with calcium trace activity and as a function of the number of cells included in the model. **(C)** Peak observed (blue) and shuffled (grey) decoding accuracy (mean, SEM) across sessions utilizing a model trained with calcium events. **(D)** Peak observed (blue) and shuffled (grey) decoding accuracy (mean, SEM) utilizing a model trained with calcium events and as a function of the number of cells included in the model.

**Supplementary Table 1.**
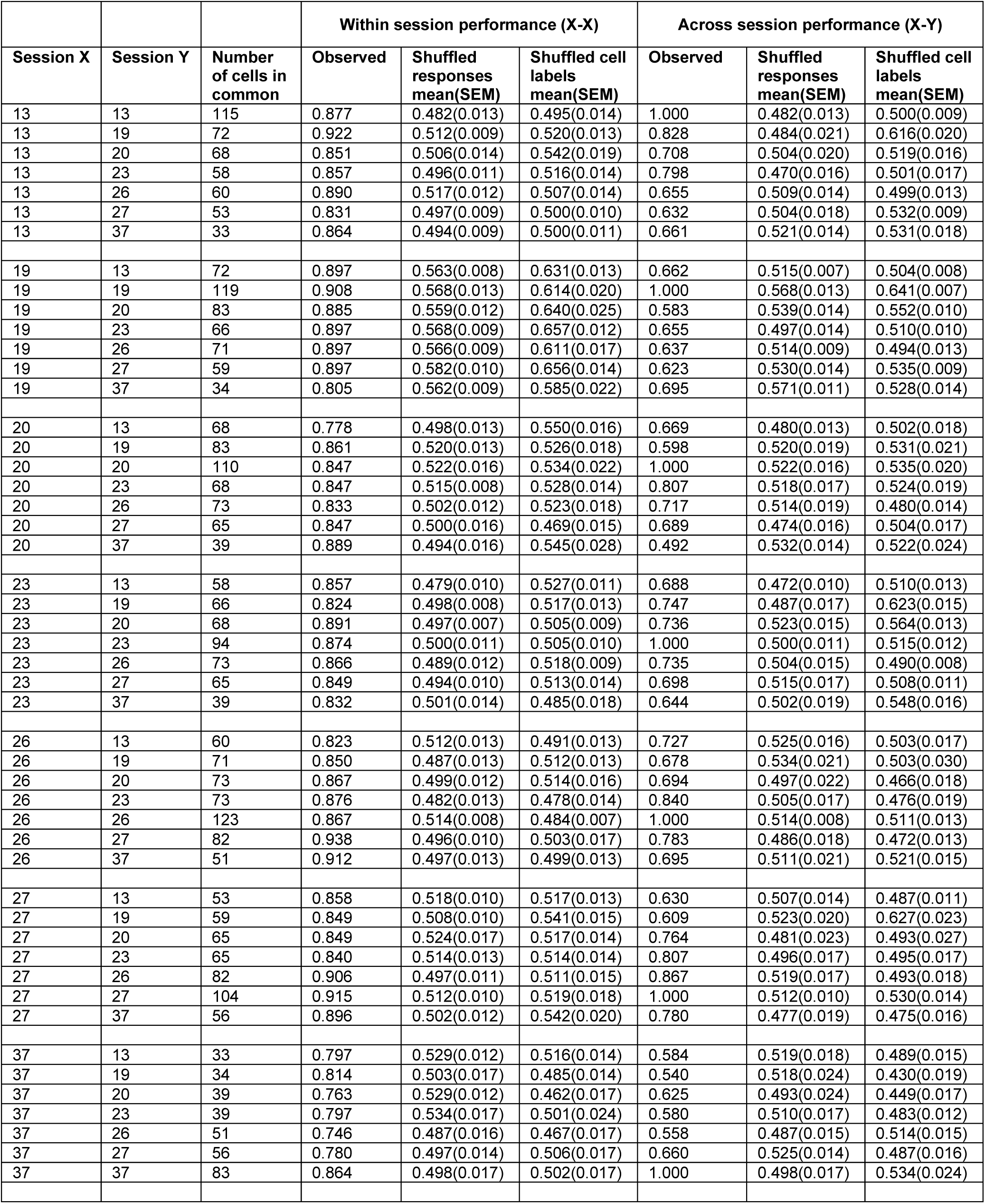
Decoding performance across sessions using longitudinally track populations of individual neurons. Within and across session decoding performance for each session pair chosen among 7 sessions spanning 24 days.

## Supplementary Videos

**Supplementary Video 1. Plug and play daily imaging with the head-mounted miniscope**. See Supplementary Material.

**Supplementary Video 2. Cellular resolution calcium imaging in dorsal premotor cortex of behaving macaque**. See Supplementary Material.

**Supplementary Video 3. Stable calcium imaging in head-unrestrained and behaving macaque**. See Supplementary Material.

**Supplementary Video 4. Sedated blood flow imaging**. See Supplementary Material.

**Supplementary Video 5. Calcium dynamics selective to contralateral arm reach direction**. See Supplementary Material.

**Supplementary Video 6. Bilateral calcium dynamics selective to left and right arm reaches**. See Supplementary Material.

## Methods

### EXPERIMENTAL MODEL AND SUBJECT DETAILS

Rhesus macaque.

The rhesus macaques were obtained from the California National Primate Research Center (CNPRC) at UC Davis as part of a Pilot Research Program awarded to the authors. The study used two healthy adult male monkeys, 5-8 years of age and weighing 13-14 kg. Animals were housed at the CNPRC at UC Davis. All experiments were conducted in compliance with the NIH Guide for the Care and Use of Laboratory Animals and were approved by the Institutional Care and Use Committee at the University of California, Davis.

### METHOD DETAILS

#### Behavioral Training

The animals were trained through positive reinforcement to exit their enclosure and enter a standard primate chair in which the animal could sit comfortably with a neck plate preventing the animal from reaching toward its head. We habituated the animal to progressively longer times sitting in the chair, as well as to soft cushions pressed lightly against the sides of the animal’s head in order to temporarily restrain the head during docking and undocking of the miniscope. The animals were then trained on a reach to reward task. For this task, the animal was seated in the primate chair at a distance of 15 cm from a tray with two predefined zones where food rewards were presented (Figure 3A, 5A). The experimenter was seated behind a panel in front of the animal and placed food rewards, through an opening in the panel, in one of the two zones chosen at random across trials. A trial began with placement of the reward followed by the animal reaching for the reward with one of the arms. A new trial began after the animal had consumed the reward from the previous trial and the arm and hand returned to a rest location on the primate chair. During a session the animal performed either a ‘single arm task’ or a ‘dual arm task’. In the single arm task, the animal was allowed to use only one arm to reach to the reward zones and the allowed arm was alternated in a block design (Figure 3A, 7A). In the dual arm task, the animal was required to use the arm directly in front of the zone, right arm to zone 1 and left arm to zone 2 (Figure 5A). Arm use was controlled by opening or closing small openings in the front panel of the primate chair. The animal was trained to perform this task for progressively longer periods of time and more trials. Only positive reinforcement was used at all stages of training.

#### Surgical Procedures

Prior to any surgeries, we obtained MR scans (3T, GE Medical Systems) from the animals in order to determine, in combination with a standard atlas, the stereotaxic coordinates for bilateral craniotomies aligned to the cortical sites (e.g. dorsal premotor cortex) targeted for virus injections and lens implantations. All of the surgical procedures that follow are depicted in schematic form (Supplementary Figure 1).

To perform virus injections, the animals were sedated and placed into a stereotaxic frame. In a first monkey (macaque 1), used only for virus testing (Figure 1B, Supplementary Figure 2), a single large craniotomy and durotomy were made over fronto-parietal cortex and multiple injections of different viruses at varying titers were made using pulled glass micropipettes (tip diameter approximately 25 μm, 1.14 mm max outer diameter) back-filled with mineral oil and connected to a microprocessor-controlled injector (NANOLITER2010; WPI, Inc.) for small volume injections. For AAV1.CaMK2a.GCaMP6f.WPRE.SV40 (UPenn Vector Core), 1 μl of fully concentrated virus (2.3E+13 GC/ml) was injected at depths of approximately 0.8-1.2 and 1.6-1.8 mm from the cortical surface for a total volume of 2 μl at each of two separate cortical sites. For the Tet-Off virus system (Salk Vector Core), AAV5.mThy1PSs.tTAad (titer: 4.35E+11 GC/ml) and AAV5.TRE3.GCaMP6f (titer: 4.16E+11 GC/ml) were first mixed together in equal volumes (1:1) for final titers for injection of half their original values and then 500 nl was injected at depths of approximately 0.8-1.2 and 1.6-1.8 mm from the cortical surface for a total volume of 1 μl at each of two separate cortical sites. Finally, we diluted with sterile PBS the 1:1 mixed viruses down to a 1:8 concentration and made similar injections at two additional cortical sites. Once all injections were complete, the dura and skull were replaced and the animal was allowed to recover and survived for 8 weeks prior to being euthanized and the brain perfused and extracted for post-mortem processing and histological examination.

In a second monkey (macaque 2), used for all of the imaging results reported in this study, we first made two craniotomies centered over dorsal premotor cortex in both the left and right hemispheres (Supplementary Figure 1). Craniotomies were approximately 10 mm in diameter with an additional 2.5 mm of thinned bone surrounding the entire perimeter of the craniotomy so that the skull would not later prevent the microendoscopic lens from reaching the desired final depth in the brain. Following completion of the craniotomies, small (approximately 3-5 mm in diameter) durotomies were made and final coordinates for virus injections were determined based on visualization of sulcal/gyral landmarks. In the right hemisphere PMd, AAV1.CaMK2a.GCaMP6f.WPRE.SV40 (UPenn Vector Core, titer: 2.3E+13 GC/ml) was injected at 2 cortical sites separated by approximately 1.5 mm in the medial-lateral axis. At each site we injected 800 nl at depths of approximately 1.2 and 2.0-2.2 mm from the cortical surface for a total volume of 1.6 μl. In the left hemisphere PMd, a 1:1 mix of AAV5.mThy1PSs.tTAad (final mixed titer: 2.18E+11 GC/ml) and AAV5.TRE3.GCaMP6f (final mixed titer: 2.08E+11 GC/ml) was injected at 2 cortical sites, again separated by approximately 1.5 mm in the medial-lateral axis. At each site we injected 500 nl at depths of approximately 1.2 and 2.0-2.2 mm from the cortical surface for a total volume of 1 μl. In all cases, injections were made at a rate of 100 nl/min and we waited approximately 10 minutes following the final injection at each site before removing the pipette from the brain. During injections, we took several photos so that we could document the location of the micropipette relative to the vessel pattern on the cortical surface. After the injections were complete, we placed four fiducial marks on the skull surrounding each of the two craniotomies to further aid in our efforts to later align lens implantation to the cortical sites injected with virus. These fiducial marks consisted of small burr-sized divots in the skull with ink placed inside and a drop of silicone elastomer (Kwik-Sil; WPI, Inc.) over the top to protect the ink from being washed away between surgeries. The four fiducial marks were placed along the ML and AP axes just beyond the perimeter of each craniotomy, aligned with the midpoint between the pair of injection sites. We measured and documented the distances from the midpoint between injections and each of these four fiducial marks. Finally, we implemented several steps to seal the durotomy and craniotomy so as to minimize neovascularization and growth of granulation tissue between surgeries. We placed artificial dura (Preclude PDX dura substitute, Gore, DE) between the cortical surface and native dura and sealed the flaps of native dura on top of the artificial dura with sterile surgical glue (Dermabond advanced, Ethicon). A second piece of artificial dura was then placed over the native dura and a thin layer of silicone elastomer was placed on top to fill in the empty volume of the craniotomy. A stainless steel metal disc (13-15 mm diameter) was placed on top of the skull, within the thinned region of bone, and sealed and secured with a combination of bone wax along its seam with the skull and a metal strap overlaying the disc and screwed to the skull. The animal’s muscle and skin were replaced and sutured to its original position over the craniotomies and the animal was allowed to recover.

Two weeks following the virus injection surgery, the animal was again sedated and placed in a stereotaxic frame for lens implantations. Once the craniotomy and durotomy were re-exposed, we used the vessel pattern on the cortical surface and measurements relative to the fiducial marks to re-identify the sites of virus injection made previously. We then made an incision in the cortex using a #11p scalpel blade (Swann-Morton, UK) approximately 2.0 mm deep and 1.2 mm across in the AP axis perpendicular to the axis of the virus injection sites. The incision was made twice, once in each direction along the AP axis, to ensure a complete incision 1.2 mm across (Kondo et al., 2018). This was immediately followed by insertion of a microendoscopic GRIN prism lens with integrated miniscope baseplate (1 mm diameter, 9 mm length Proview Integrated Prism Lens; Inscopix, Inc.) aligned to the incision site. The prism probe was inserted at a rate of approximately 500 μm/min. to a final depth of approximately 2.0 mm from the cortical surface. Once the lens was at its final depth, we placed a layer of silicone elastomer over the exposed brain surrounding the lens. We then placed a metal disc (13-15 mm diameter) over the craniotomy, with a 2 mm wide slot running half its diameter to allow for the lens to pass through. We sealed the slot in the metal disc with silicone elastomer and then proceeded to chronically secure the lens to the skull by placing around the lens an aluminum cranial chamber (3 mm height, 17 mm diameter) and embedding the lens within a combination of cement (Metabond, Parkell, NY) and acrylic inside the cranial chamber. The cranial chamber was chronically secured to the skull with a combination of acrylic and several skull screws resulting in a single, large acrylic headcap securing both chambers bilaterally. A removable chamber cap was placed on each chamber to protect the miniscope baseplate and lens from debris or damage between imaging sessions. The animal was allowed to recover for two weeks prior to the first calcium imaging session.

#### Imaging during Behavioral Sessions

Calcium imaging signals were acquired using the nVista system (Inscopix, Inc., Palo Alto, CA), which includes a miniscope (nVista 3.0) that docks to the baseplate-integrated lens implanted in the brain (Figure 1A). The signals are transferred by a thin, flexible cable to the data acquisition system (DAQ) where they are recorded and saved. Imaging parameters, including frame rate, LED power, sensor gain and electronic focus were all controlled using the Inscopix Data Acquisition Software (IDAS; Inscopix, Inc.) running in a web browser (Chrome, Firefox) on a computer that communicated wirelessly with the DAQ. The animal’s motor reach behavior was monitored by overhead camera (Allied Vision, Stingray F-033C) and recorded using the CinePlex system (Plexon Inc., Dallas, Texas) controlled by CinePlex software running on a separate, dedicated computer. The calcium imaging and behavior monitoring were synchronized by sending TTLs corresponding to each nVista frame capture, from the imaging system to the behavioral monitoring system. Additionally, recordings on the behavioral monitoring system were triggered by the Inscopix imaging system, again via TTL. When performing multisite imaging simultaneously from both hemispheres, a second nVista 3.0 system was used that also sent synchronizing TTLs to the behavioral monitoring system.

Daily sessions combining imaging and behavior began by first positioning the animal seated in the primate chair in the experimental setup (described above). The animal’s head was temporarily (approximately 5 minutes) restrained with soft cushions pressed against the sides of the head to allow for removal of the chamber cap and miniscope baseplate cover followed by docking and securement (by tightening a single set screw) of the miniscope to the baseplate (Figure 1A, Supplementary Video 1). The animal’s hands were tracked in real-time using the CinePlex ‘color tracking’ mode, and the top of the animal’s hands were shaved and painted with colored (pink or purple) ink for this purpose. During a session the animal performed either the ‘single arm’ task or the ‘dual arm’ task (see above). During a typical session the animal performed approximately 250 trials, and data collection lasted about one hour. At the end of data collection, the animal’s head was again temporarily restrained to undock the miniscopes and replace the cap on the chamber. Calcium imaging signals were acquired at 20 Hz (LED power 0.4; Gain 5) in the left hemisphere and at 10 Hz (LED power 0.8; Gain 7) in the right hemisphere.

#### Doxycycline Administration and Testing

To study how Doxycycline (Dox) regulates transgene (i.e. GCaMP) expression of the AAV Tet-Off system, Dox was administered (15 mg/kg; 5 days) orally by mixing it with food. Calcium imaging signals were acquired prior, during and up to 6 weeks after cessation of Dox administration to quantify changes in calcium imaging performance (fluorescence levels, cell numbers, calcium event kinetics) thought to correlate with levels of GCaMP expression.

#### Sedated blood flow imaging

The one-photon miniscope has been used previously to perform in vivo blood flow imaging in rodents (Ghosh et al., 2011). To study the feasibility of blood flow imaging in the macaque brain the animal was sedated and administered Fluorescein (AK-Fluor 10%, Akorn Inc., IL) intravenously through an IV line placed in the arm. 1-2 ml Fluorescein was injected at a rate of 1 ml/min either manually or with the aid of an injection pump. Additionally, a slow drip (0.1 ml/min) of Fluorescein was given to maintain stable fluorescence intensity levels during the imaging session. Imaging signals were acquired starting a few seconds before Fluorescein administration and lasted approximately 15 mins (Supplementary Video 4).

#### Post-mortem assessment

For post-mortem assessment, animals were euthanized and transcardially perfused with saline, and then a series of 4% paraformaldehyde alone, followed by 10% and 20% sucrose in 4% paraformaldehyde. The brains were carefully extracted and soaked in 30% sucrose in 4% paraformaldehyde for 24 hours prior to being placed in 30% sucrose in PBS for several days. Once the brain was sufficiently cryoprotected with sucrose, we cut 50 μm thick coronal sections of the brain on a freezing microtome. Sections that included the injection sites and lens implantation sites were stained for cell nuclei (DAPI), mounted on slides and cover-slipped for conventional benchtop fluorescence microscopy. GCaMP expression patterns and expression levels were assessed based on native GCaMP fluorescence. No antibody staining was conducted.

To compare GCaMP expression levels across cases, we used ImageJ to draw regions of interest (ROIs) and calculate the integrated fluorescence density within these ROIs. These values were normalized by the area of each ROI and the exposure time used to capture the image. For each virus injection site of either the Tet-Off virus system or the conventional CaMK2a virus we made these measurements with an ROI centered within the region of peak expression and subtracted from that a second ROI centered within a region without expression (background fluorescence within the same tissue section). An average was taken across the two separate virus injection sites tested for each virus.

### QUANTIFICATION AND STATISTICAL ANALYSIS

#### Behavioral tracking

CinePlex Editor software (Plexon Inc, TX) was used to track the reach trajectories offline and to identify the times when the animal’s hand entered the reward zones. Although the animal was required to bring the hand to a ‘rest’ position before a trial began, the orientation of the hand was often variable, which made it difficult to reliably identify when the hand was in the ‘rest’ position. We therefore used the timing of reward zone entries, which were more reliably identified, to study the reach related modulation of the calcium imaging signals.

#### Image processing

The calcium imaging videos were processed using a custom pipeline developed using the Inscopix Data Processing Software (IDPS; Inscopix, Inc.) application programming interface (API). The processing pipeline includes a first step that corrects defective pixels, appropriately spatially crops the FOV, and spatially downsamples (4X) each of the imaging frames to reduce the size of the data while maintaining acceptable resolution and SNR. The video frames were then spatially filtered (band-pass) with cut-offs set to 0.005 pixel^-1^ (low) and 0.5 pixel^-1^ (high). Following spatial filtering the imaging frames were corrected for brain movement artifacts associated with respiration and head-unrestrained behavior using a standard rigid translation algorithm (Thévenaz et al., 1998). Cell segmentation and extraction of calcium activity traces of individual cells was performed by using the CNMFe algorithm (Zhou et al., 2018) on the motion-corrected imaging frames. The observed imaging data is modeled as an additive superposition of all cells’ spatiotemporal activity, time-varying background and noise. Each cell is characterized by its spatial footprint characterizing the cell’s shape and location. Segmented cells were manually curated and cells with discontinuous and non-cell-like shapes or cells with activity traces with non-calcium like events (e.g. blood vessels falsely identified as cells) were discarded. Accepted cells and their activity traces were exported to Matlab (MathWorks, MA) for further analysis.

#### Analysis Head movement and motion correction

The motion correction algorithm (Thévenaz et al., 1998) applies to each frame a translation that minimizes the difference between the transformed frame and the reference frame. The ‘mean frame’, which is the average of all frames following spatial filtering, was used as the reference frame. The median of the applied motion correction translations for each frame was taken as an estimate of the movement induced noise during imaging. The efficacy of the motion correction algorithm in mitigating the animal movement-induced motion artifacts in the imaging data was evaluated by estimating the correlation between the head movement and the average signal intensity of an ROI defined at the edge of a large blood vessel in the imaging FOV (Figure 1C). Head movement was estimated offline by tracking one of the chamber caps or the miniscope using the CinePlex Editor software (Plexon Inc, TX) as described above. Average signal intensity in a 34×50 µm ROI was estimated for each frame before and after applying motion correction and correlated with head movement using Pearson linear correlation.

#### Cell maps and basic cell metrics

Cell maps containing spatial footprints of all cells were generated as the average of the normalized cell images of individual cell footprints. Average fluorescence across the FOV was estimated after rectangularly cropping the raw imaging frames to exclude pixels outside the perimeter of the lens. Calcium events were determined from the CNMFe extracted (dF) calcium activity traces for each cell using a custom algorithm in the IDPS API. Briefly, the algorithm first identifies time points where the slope of the calcium activity trace changes from positive to negative (p->n) and from negative to positive (n->p) resembling a monotonic rise followed by an exponentially decaying calcium event. For all of the p->n points, it then takes the activity value of the previous n->p plus the scaled median absolute deviation (MAD) to calculate a dynamic activity threshold curve. The scaling parameter is set by the event minimal size parameter (set to 4 here). Events are detected whenever the calcium activity time trace goes above the dynamic event detection threshold. The final rising event times are determined after removing any event times for which the decay after the entered tau-off parameter (set here to 0.2 s) is greater than expected assuming a single-exponential decay model. Rise and decay times and SNR are estimated around each identified event and summarized for each cell by taking the median value.

#### Reach related calcium activity analysis

Rasters of calcium events were constructed by aligning them around the time of reach entry into reward zones. Peri-event histograms were constructed by estimating the average number of events in 1 s sliding window with a step size of 0.5 s. Heatmaps showing task related modulation of population calcium activity were constructed after z-scoring the average peri-event calcium activity traces. Cells in the heatmaps were sorted by their selectivity to reach direction or hand used. Cells were characterized as reach-modulated cells if the activity during reach was significantly modulated during the reach epoch ([0 1]s interval with respect to reward zone entry) compared to a baseline epoch ([-3 -2]s interval with respect to reward zone entry). The average of the calcium activity trace in the reach and baseline epochs were used as signal and noise respectively to compute the area under the receiver operator characteristic curve (AUC). Bootstrap resampling was used to estimate 95% confidence intervals. Reach direction selectivity of each cell was characterized by estimating a tuning index (TI), TI = (Ca^Z1^ - Ca^Z2^)/(|Ca^Z1^| + |Ca^Z2^|) where Ca^Z1^ and Ca^Z2^ are the average calcium activity traces in [0 1]s intervals with respect to entry into reward zones 1 and 2 respectively. Bootstrap resampling was used to estimate 95% confidence intervals.

To investigate whether cells that were similarly selective were spatially clustered together, we estimated among the reach modulated cells the frequency with which the closest neighboring cell was similarly selective. For significance testing this estimate was compared to a null distribution generated by multiple (100) random shuffles of the cell labels. To investigate whether the prevalence of reach direction selectivity depended on the depth in cortex, we estimated, among the reach modulated cells, the proportion of cells that were selective as a function of cortical depth using 75 µm windows with step size of 3 µm.

#### Decoding

Reach related modulation of calcium activity traces allowed for decoding of the animal’s behavior on individual trials. Decoding accuracy was computed as a function of time ([-2 2]s) around the time of reward zone entry. Partial Least Squares Discriminant Analysis (PLS-DA) with leave one out cross-validation was used to predict the animal’s reach direction using calcium activity traces in a 400 ms sliding window, and a 100 ms step size, around the time of reach. 95% confidence intervals and standard errors were computed from Monte Carlo bootstrap distributions simulated from R=50 random draws with replacement from sample datasets. Reach direction could also be decoded with discrete times of calcium events instead of the continuous activity traces. Logistic regression with 10-fold cross-validation was used on population activity in a 2 s sliding window, and a 100 ms step size. Chance level decoding accuracy was characterized by repeating the decoding analysis after shuffling the reach direction labels across trials. Peak decoding accuracy within the [-2 2]s window around the time of reach was noted to plot decoding accuracy across sessions. Dependence of decoding accuracy on the number of cells was investigated by averaging the peak decoding accuracy estimated during multiple random subsamples of cells.

#### Longitudinal registration

We registered cells across multiple sessions based on the similarity in their spatial footprints by using a custom algorithm in IDPS that is similar to the one described in Sheintuch et. al (2017). The registration is performed in two steps: cell map alignment and cell matching. Specifically, for each session, we generated a cell map as the average of the normalized cell images of individual cell footprints (*S*_*Pc*_) extracted using CNMFe

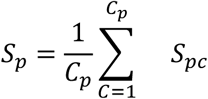

where *C*_*P*_ is the number of cells. We then aligned each cell map (*S*_*P*_) to a reference map (*S*_*r*_) using the enhanced correlation coefficient (*ECC*) image registration algorithm (Evangelidis and Psarakis, 2008), a rigid transformation that allows for translation and rotation. When registering cells between two sessions (e.g. Figures 2D, Figure 4, Supplementary Figure 6A,B) one of the sessions acted as the reference map. The cell map from the session closest to the midpoint between the first and last session was used as the reference map when tracking the same cells through more than two sessions (e.g. Figure 2F, Supplementary Figure 6C). The rectangular region for which all of the aligned cell maps overlapped was computed and defined as the common and registered FOV (*R*) for all sessions. The estimated transformation (*T*_*P*_) and overlap region of each cell map was applied to the individual cell footprints *S*_*Pc*_, resulting in registered footprints 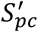.

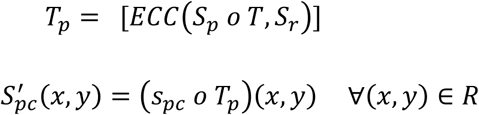

Registered footprints all have the same number of pixels, and the registered footprint of the same cell in different sessions should mostly overlap. Some cells may only be active in particular sessions and, therefore, would not overlap (match) with any cell footprint in other sessions. In order to find all unique cells (with or without matching cells) present in this longitudinal study, we defined a global cell set that is initialized to be all of the cells in the reference session, and then incrementally modified this global cell set by finding matches between it and other aligned cell sets. For each cell in an aligned cell set, we chose the cell in the current global cell set that had the highest normalized cross correlation (*NCC*). If the highest *NCC* passed a set threshold, it was identified as a match to the existing global cell, and we updated the global cell footprint as the mean of all matched footprints; otherwise, the cell was treated as a new cell added into the global cell set. The matching process was repeated until there were no possible matches with a correlation above the set threshold. We profiled the matching results with a series of *NCC* thresholds, and reported results for 0.3, 0.5 and 0.7.

#### Longitudinal relationship between calcium activity and behavior

Calcium imaging allows for tracking of a subset of cells across sessions and days. Changes in reach direction selectivity across sessions was characterized by evaluating correlation between tuning indices and the difference in tuning indices for cells tracked between a pair of sessions. Similarly, tracking the cells across sessions allowed for prediction of behavior during a (test) session by using a decoding model trained on a different (training) session, showing that decoding model training can transfer between sessions. Within session decoding was performed by leave-one-out cross-validation as described above. Across session decoding was performed using a decoding model trained on all trials in the training session. Chance level decoding accuracy was estimated by repeating the decoding analysis after randomly shuffling the labels of the longitudinally tracked cells between sessions. The transfer of decoding model training was further characterized by taking the ratio of decoding accuracy during the test session and the training session and plotting this ratio as a function of the intersession interval (Figure 4E).

### DATA AND CODE AVAILABILITY

Datasets and custom code generated during the study are available from the corresponding author upon request.

## Supplementary Material

All supplementary materials can be found online.

Supplementary Figures 1-8 and Table 1.

Supplementary Videos 1-6.

## Author Contributions

J.N., J.C., K.M. and J.M. conceived the project. J.N., A.B., S.S., J.C. and K.M. designed the experiments. J.N., A.B., S.S. and J.C. performed the surgeries. A.B. and R.E performed the imaging and behavior experimental sessions. J.N., A.B., S.S., R.E. and S.X. analyzed the data. All authors contributed to writing the paper.

## Acknowledgements

This research was funded by a grant from the California National Primate Research Center (CNPRC) Pilot Research Program. We are grateful to Morrison lab member Sean Ott for virus preparation support and the CNPRC staff for support with behavioral training, surgery, pathology and general care of animals. We thank Inscopix staff, including Arash Tajik, Sam Malanowski, Evan DeJarnette and Mark Trulson, for supporting design, manufacturing and testing of custom imaging components and David Cheng for testing of viruses. We thank Moxon lab members, Ashley Schnider and Megan Santos, for support with offline tracking of reaching behavior. We appreciate the helpful comments on the manuscript form Drs. Krishna Shenoy, Alice Stamatakis, Mark Miller and Shay Neufeld.

## Notes

### Competing Interest Statement

Anil Bollimunta, Pei Xu and Jonathan Nassi are paid employees of Inscopix, Inc.

