## Supplementary Figures and Table for "Head-mounted microendoscopic calcium imaging in dorsal premotor cortex of behaving rhesus macaque"

**Supplementary Material**

Anil Bollimunta[1*], Samantha R. Santacruz[2*], Ryan W. Eaton[3,4], Pei S. Xu[1], John H. Morrison[3,4], Karen A. Moxon[3,4], Jose M. Carmena[5**], Jonathan J. Nassi[1**°]

[1]Inscopix, Inc.; Palo Alto, CA, USA
[2]University of Texas at Austin; Austin, TX, USA
[3]California National Primate Research Center; Davis, CA, USA
[4]University of California, Davis; Davis, CA, USA
[5]University of California, Berkeley; Berkeley, CA, USA

*These authors contributed equally to this work
**Senior authors
°Corresponding author and lead contact

### Supplementary Figures and Table

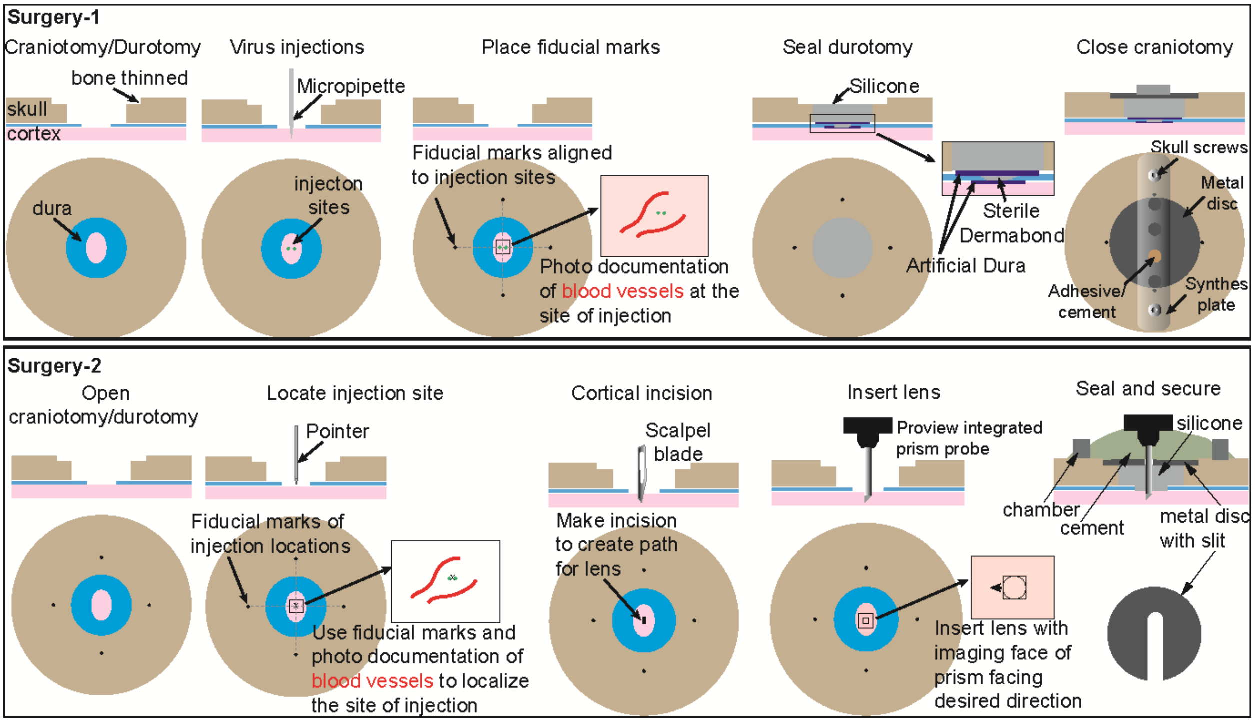

**Supplementary Figure 1. Schematic of surgical steps to prepare macaque for chronic imaging.**

Top: A schematic of surgical steps during the first surgery for injections of virus.

Bottom: A schematic of surgical steps during the second surgery for lens implantation.

See Methods for more details.

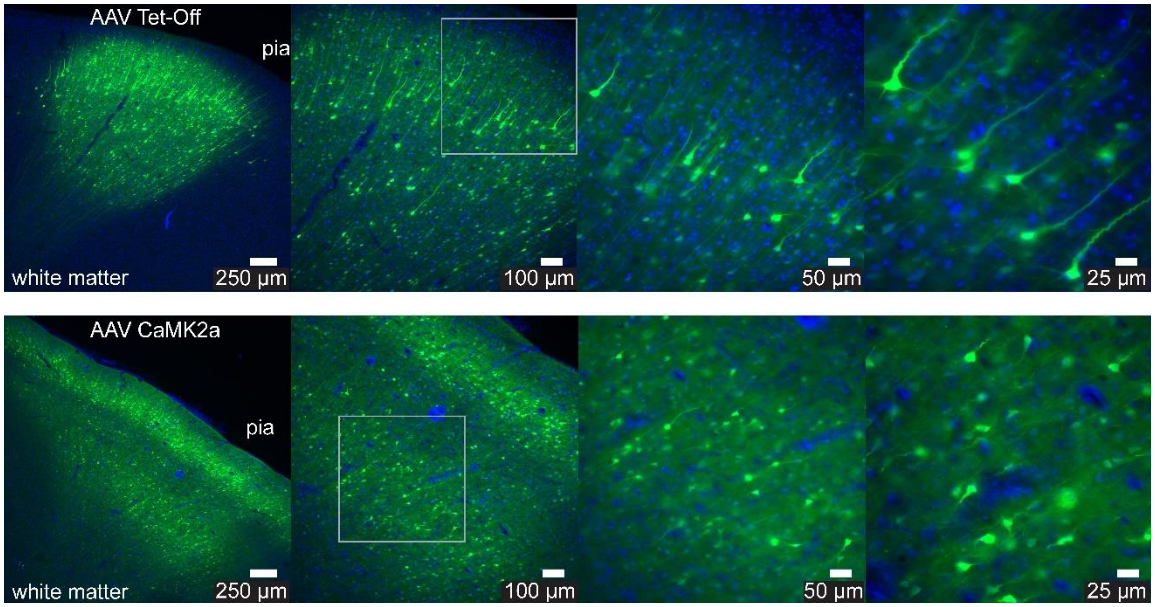

**Supplementary Figure 2. Post-mortem assessment of GCaMP expression in macaque 1.** Top: Post-mortem native GCaMP expression (green) and DAPI-stained cell nuclei (blue) in the cortex 8 weeks following injections of the AAV Tet-Off virus system (top) or AAV1 CaMK2a virus (bottom) in animal 1. Images of increasing magnification are shown from left to right, with scale bars equal to 250, 100, 50 and 25 μm respectively. The higher magnification images to the right are from the superficial (top row) or deep (bottom row) layers indicated by the rectangle in the middle-left panel.

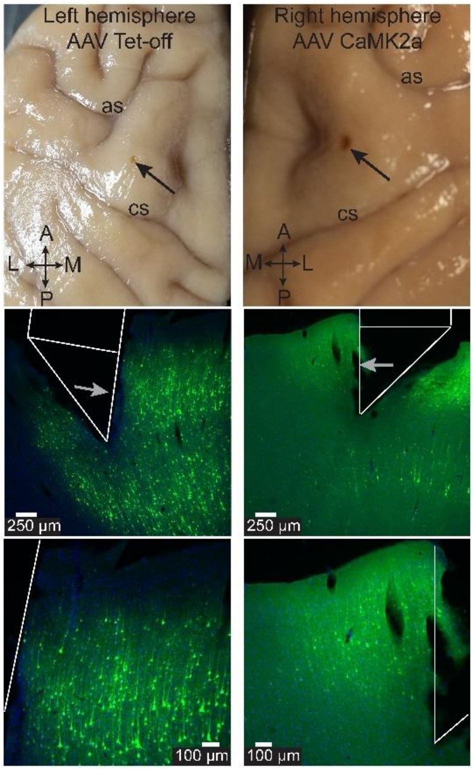

**Supplementary Figure 3. Post-mortem assessment of GCaMP expression and location of lens implants in macaque 2.**

Top: Post-mortem, ex vivo image of dorsal surface of brain and lens implant site for the left hemisphere (left; AAV Tet-Off) and right hemisphere (right; AAV CaMK2a) from animal 2. The arrow indicates the location of the lens implant. ‘as’ arcuate sulcus, ‘cs’ central sulcus. ‘A’ anterior, ‘P’ posterior, ‘M’ medial, ‘L’ lateral. Middle: Post-mortem native GCaMP expression (green) and DAPI-stained cell nuclei (blue) in PMd cortex 8.5 months following injections of the AAV Tet-Off virus system (left) or AAV CaMK2a (right) in animal 2. The estimated location of the prism lens during imaging is outlined in white. The arrow indicates the imaging surface of the prism and the direction of imaging. M-L axis as in the top panels. Scale bars equal 250 μm. Bottom: Same as above at higher magnification. Scale bars equal 100 μm.

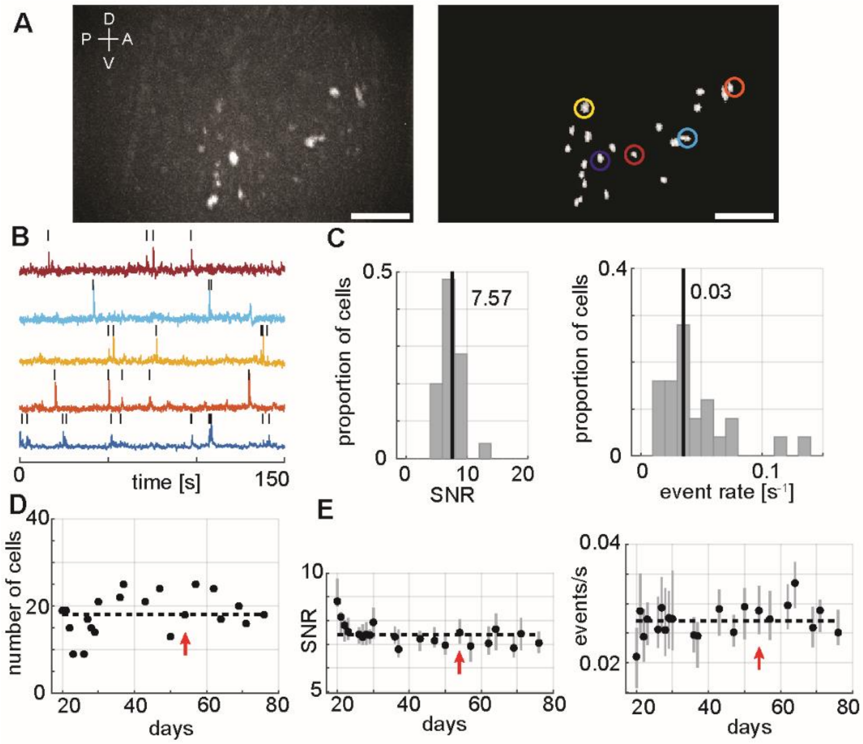

**Supplementary Figure 4. Cellular resolution imaging in macaque dorsal premotor cortex right hemisphere.**

**(A)** Left: Max projection image of in vivo GCaMP fluorescence in the right hemisphere PMd over the course of a single example session. The bright colored regions in the image indicate cells that exhibited active calcium dynamics during the recording. Dorsal (D), Ventral (V), Anterior (A), Posterior (P) denote orientation in the premotor cortex. Scale bar equals 250 μm. Right: Map of cells extracted using CNMFe from the same example session. Colored circles indicate example cell calcium activity traces in (B). Scale bar equals 250 μm. **(B)** Calcium activity (dF, peak normalized) traces of example cells highlighted in (A). The black tick marks above the traces indicate CNMFe extracted calcium events. **(C)** Distribution of median calcium event SNR (left) and median calcium event rate (right) for the entire population of cells recorded in the example session. The vertical lines indicate the median SNR (7.57) and event rate (0.03) values. **(D)** Number of cells that were imaged for each session across 60 days. The dashed line indicates the mean value. The red arrow indicates the example session. **(E)** Calcium event SNR (left) and rates (right) (median and IQR) for each session across 60 days. The dashed line indicates the mean value. The red arrow indicates the example session.

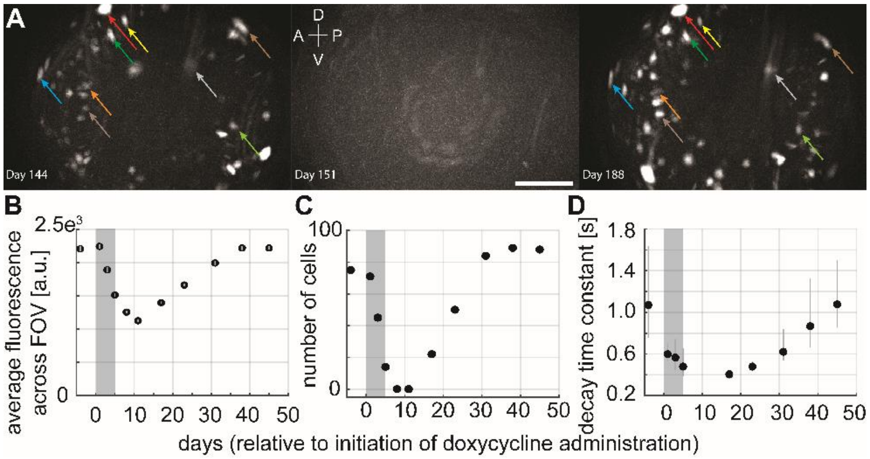

**Supplementary Figure 5. Control of Tet-Off virus mediated GCaMP expression with Doxycycline.**

**(A)** Max projection images of in vivo GCaMP fluorescence 1 day prior to initiation of Doxycycline (Dox) administration (left), 8 days following initiation (3 days following end) of Dox administration (middle) and 45 days following initiation (40 days following end) of Dox administration (right). Colored arrows indicate putative same-cells pairs identified 1 day prior to and 45 days following the initiation of Dox administration. Dorsal (D), Ventral (V), Anterior (A), Posterior (P) denote orientation in the premotor cortex. Scale bar equals 250 μm. **(B)** Average fluorescence across the FOV before, during and after Dox administration (mean, SEM). Grey shaded region indicates the time period of Dox administration. Fluorescence levels were reduced following Dox administration and returned to baseline approximately 35 days following its cessation. **(C)** Number of cells that were imaged for each session before, during and after Dox administration. Grey shaded region indicates the time period of Dox administration. The number of cells was reduced following Dox administration and returned to baseline approximately 30 days following its cessation. **(D)** Calcium event decay time constants (mean, 95% CI) before, during and after Dox administration. Grey shaded region indicates the time period of Dox administration. Decay time constants were reduced following Dox administration and returned to baseline approximately 40 days following its cessation.

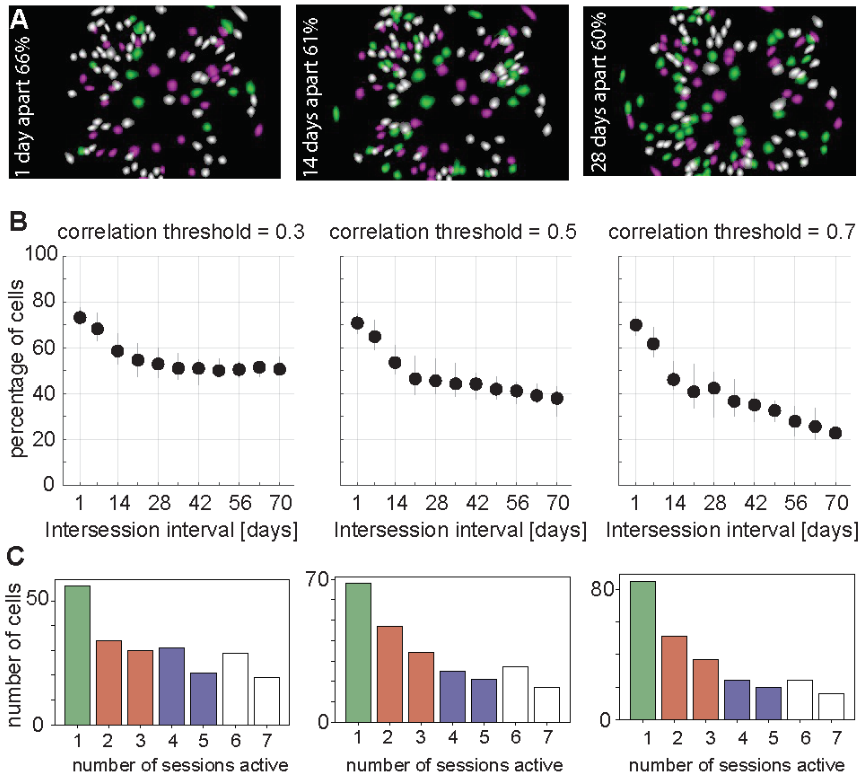

**Supplementary Figure 6. Dependencies of longitudinal tracking performance across sessions.**

**(A)** Overlays of CNMFe-extracted cell maps from two separate sessions (magenta and green cells) spaced 1 (left), 14 (middle) and 28 (right) days apart. The percentage of cells present and active in both sessions (white cells) is reported in each case. **(B)** Percentage of cells (median, IQR) in common between two sessions as a function of the intersession interval (days) for different spatial correlation thresholds used to determine putative 'same-cell' pairs (left = 0.3, middle = 0.5, right = 0.7). **(C)** Percentage of cells as a function of the number of sessions (non-consecutive) found to be present and active for different spatial correlation thresholds (left = 0.3, middle = 0.5, right = 0.7).

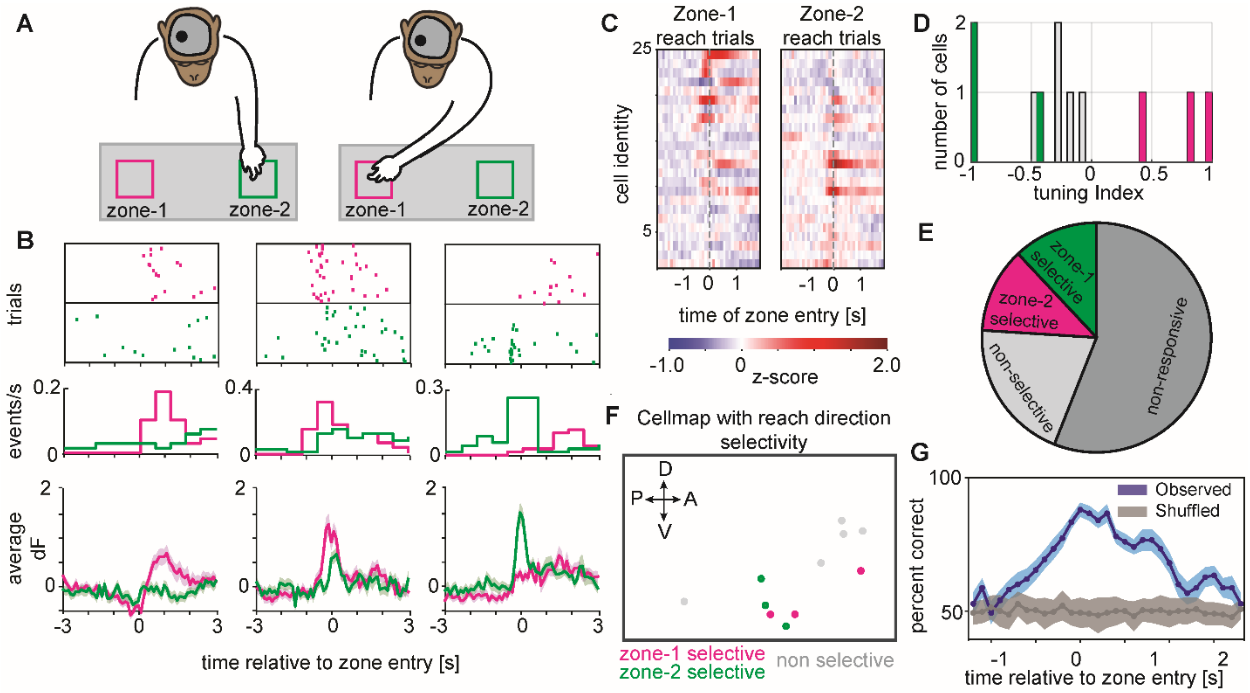

**Supplementary Figure 7. Direction selective calcium dynamics in right hemisphere PMd and decoding of motor reach behavior.**

**(A)** Schematic of the macaque performing the reach to reward task with an nVista miniscope mounted on the head to image from right hemisphere PMd. In these sessions, the macaque reached with the left arm (contralateral to the imaged hemisphere) to one of two zones, either zone 1 (magenta) or zone 2 (green). **(B)** Three example cells from the right hemisphere PMd exhibiting zone 1 selectivity (left), zone 2 selectivity (right) or nonselective modulation to either reach location (middle) in a single example session. Top: Rasters of calcium event times across multiple trials aligned to the time of reach entry (dashed vertical line) into zone 1 (magenta) or zone 2 (green). Middle: Peri-stimulus time histogram (PSTH) of calcium events as a function of time relative to reach entry into zone 1 (magenta) or zone 2 (green). Bottom: Calcium trace activity (mean, SEM) as a function of reach entry into zone 1 (magenta) or zone 2 (green). **(C)** Heatmap depicting z-scored trial-averaged calcium trace activity for each cell in the population (rows) on either zone 1 (left) or zone 2 (right) reach trials and aligned to the time of zone entry (dashed vertical line). The cells have been sorted top to bottom based on their selectivity (tuning index) to zone 1 or zone 2 reaches respectively. **(D)** Top: Distribution of reach direction selectivity (tuning index) for the entire population of cells recorded in the example session. Magenta and green colored bars indicate cells that had significant ($p < 0.05$; see Methods) reach direction selectivity to zone 1 (positive tuning indices) or zone 2 (negative tuning indices) respectively. **(E)** Pie chart depicting the percentage of cells in the example session that were classified as zone 1 selective (magenta), zone 2 selective (green), reach modulated but nonselective (light grey) or nonresponsive (dark grey). **(F)** Cell map depicting the spatial distribution of reach direction selectivity. Cells selective for zone 1 (magenta), zone 2 (green) or reach modulated but nonselective (light grey) are indicated. Dorsal (D), Ventral (V), Anterior (A), Posterior (P) denote orientation in the premotor cortex. **(G)** Observed accuracy of decoding the animal's reach direction on individual trials (mean, SEM) utilizing a model trained with calcium trace activity in 400 ms time bins (and 100 ms steps) around the time of reach entry into zones 1 and 2 (blue). Chance level decoding accuracy estimated by shuffling the reach direction across trials (grey).

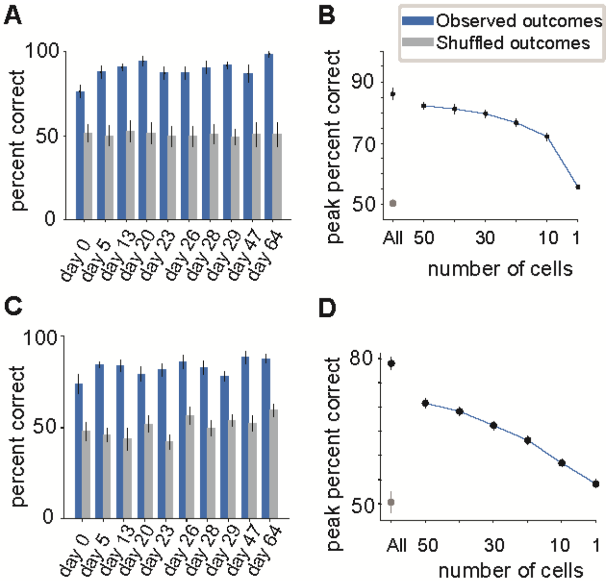

**Supplementary Figure 8. Decoding of motor reach behavior with calcium activity traces and events.**

**(A)** Peak observed (blue) and shuffled (grey) decoding accuracy (mean, SEM) across sessions utilizing a model trained with calcium trace activity. **(B)** Peak observed (blue) and shuffled (grey) decoding accuracy (mean, SEM) utilizing a model trained with calcium trace activity and as a function of the number of cells included in the model. **(C)** Peak observed (blue) and shuffled (grey) decoding accuracy (mean, SEM) across sessions utilizing a model trained with calcium events. **(D)** Peak observed (blue) and shuffled (grey) decoding accuracy (mean, SEM) utilizing a model trained with calcium events and as a function of the number of cells included in the model.

| | | | Within session performance (X-X) | | | Across session performance (X-Y) | | |
|---|---|---|---|---|---|---|---|---|
| **Session X** | **Session Y** | **Number of cells in common** | **Observed** | **Shuffled responses mean(SEM)** | **Shuffled cell labels mean(SEM)** | **Observed** | **Shuffled responses mean(SEM)** | **Shuffled cell labels mean(SEM)** |
| 13 | 13 | 115 | 0.877 | 0.482(0.013) | 0.495(0.014) | 1.000 | 0.482(0.013) | 0.500(0.009) |
| 13 | 19 | 72 | 0.922 | 0.512(0.009) | 0.520(0.013) | 0.828 | 0.484(0.021) | 0.616(0.020) |
| 13 | 20 | 68 | 0.851 | 0.506(0.014) | 0.542(0.019) | 0.708 | 0.504(0.020) | 0.519(0.016) |
| 13 | 23 | 58 | 0.857 | 0.496(0.011) | 0.516(0.014) | 0.798 | 0.470(0.016) | 0.501(0.017) |
| 13 | 26 | 60 | 0.890 | 0.517(0.012) | 0.507(0.014) | 0.655 | 0.509(0.014) | 0.499(0.013) |
| 13 | 27 | 53 | 0.831 | 0.497(0.009) | 0.500(0.010) | 0.632 | 0.504(0.018) | 0.532(0.009) |
| 13 | 37 | 33 | 0.864 | 0.494(0.009) | 0.500(0.011) | 0.661 | 0.521(0.014) | 0.531(0.018) |
| 19 | 13 | 72 | 0.897 | 0.563(0.008) | 0.631(0.013) | 0.662 | 0.515(0.007) | 0.504(0.008) |
| 19 | 19 | 119 | 0.908 | 0.568(0.013) | 0.614(0.020) | 1.000 | 0.568(0.013) | 0.641(0.007) |
| 19 | 20 | 83 | 0.885 | 0.559(0.012) | 0.640(0.025) | 0.583 | 0.539(0.014) | 0.552(0.010) |
| 19 | 23 | 66 | 0.897 | 0.568(0.009) | 0.657(0.012) | 0.655 | 0.497(0.014) | 0.510(0.010) |
| 19 | 26 | 71 | 0.897 | 0.566(0.009) | 0.611(0.017) | 0.637 | 0.514(0.009) | 0.494(0.013) |
| 19 | 27 | 59 | 0.897 | 0.582(0.010) | 0.656(0.014) | 0.623 | 0.530(0.014) | 0.535(0.009) |
| 19 | 37 | 34 | 0.805 | 0.562(0.009) | 0.585(0.022) | 0.695 | 0.571(0.011) | 0.528(0.014) |
| 20 | 13 | 68 | 0.778 | 0.498(0.013) | 0.550(0.016) | 0.669 | 0.480(0.013) | 0.502(0.018) |
| 20 | 19 | 83 | 0.861 | 0.520(0.013) | 0.526(0.018) | 0.598 | 0.520(0.019) | 0.531(0.021) |
| 20 | 20 | 110 | 0.847 | 0.522(0.016) | 0.534(0.022) | 1.000 | 0.522(0.016) | 0.535(0.020) |
| 20 | 23 | 68 | 0.847 | 0.515(0.008) | 0.528(0.014) | 0.807 | 0.518(0.017) | 0.524(0.019) |
| 20 | 26 | 73 | 0.833 | 0.502(0.012) | 0.523(0.018) | 0.717 | 0.514(0.019) | 0.480(0.014) |
| 20 | 27 | 65 | 0.847 | 0.500(0.016) | 0.469(0.015) | 0.689 | 0.474(0.016) | 0.504(0.017) |
| 20 | 37 | 39 | 0.889 | 0.494(0.016) | 0.545(0.028) | 0.492 | 0.532(0.014) | 0.522(0.024) |
| 23 | 13 | 58 | 0.857 | 0.479(0.010) | 0.527(0.011) | 0.688 | 0.472(0.010) | 0.510(0.013) |
| 23 | 19 | 66 | 0.824 | 0.498(0.008) | 0.517(0.013) | 0.747 | 0.487(0.017) | 0.623(0.015) |
| 23 | 20 | 68 | 0.891 | 0.497(0.007) | 0.505(0.009) | 0.736 | 0.523(0.015) | 0.564(0.013) |
| 23 | 23 | 94 | 0.874 | 0.500(0.011) | 0.505(0.010) | 1.000 | 0.500(0.011) | 0.515(0.012) |
| 23 | 26 | 73 | 0.866 | 0.489(0.012) | 0.518(0.009) | 0.735 | 0.504(0.015) | 0.490(0.008) |
| 23 | 27 | 65 | 0.849 | 0.494(0.010) | 0.513(0.014) | 0.698 | 0.515(0.017) | 0.508(0.011) |
| 23 | 37 | 39 | 0.832 | 0.501(0.014) | 0.485(0.018) | 0.644 | 0.502(0.019) | 0.548(0.016) |
| 26 | 13 | 60 | 0.823 | 0.512(0.013) | 0.491(0.013) | 0.727 | 0.525(0.016) | 0.503(0.017) |
| 26 | 19 | 71 | 0.850 | 0.487(0.013) | 0.512(0.013) | 0.678 | 0.534(0.021) | 0.503(0.030) |
| 26 | 20 | 73 | 0.867 | 0.499(0.012) | 0.514(0.016) | 0.694 | 0.497(0.022) | 0.466(0.018) |
| 26 | 23 | 73 | 0.876 | 0.482(0.013) | 0.478(0.014) | 0.840 | 0.505(0.017) | 0.476(0.019) |
| 26 | 26 | 123 | 0.867 | 0.514(0.008) | 0.484(0.007) | 1.000 | 0.514(0.008) | 0.511(0.013) |
| 26 | 27 | 82 | 0.938 | 0.496(0.010) | 0.503(0.017) | 0.783 | 0.486(0.018) | 0.472(0.013) |
| 26 | 37 | 51 | 0.912 | 0.497(0.013) | 0.499(0.013) | 0.695 | 0.511(0.021) | 0.521(0.015) |
| 27 | 13 | 53 | 0.858 | 0.518(0.010) | 0.517(0.013) | 0.630 | 0.507(0.014) | 0.487(0.011) |
| 27 | 19 | 59 | 0.849 | 0.508(0.010) | 0.541(0.015) | 0.609 | 0.523(0.020) | 0.627(0.023) |
| 27 | 20 | 65 | 0.849 | 0.524(0.017) | 0.517(0.014) | 0.764 | 0.481(0.023) | 0.493(0.027) |
| 27 | 23 | 65 | 0.840 | 0.514(0.013) | 0.514(0.014) | 0.807 | 0.496(0.017) | 0.495(0.017) |
| 27 | 26 | 82 | 0.906 | 0.497(0.011) | 0.511(0.015) | 0.867 | 0.519(0.017) | 0.493(0.018) |
| 27 | 27 | 104 | 0.915 | 0.512(0.010) | 0.519(0.018) | 1.000 | 0.512(0.010) | 0.530(0.014) |
| 27 | 37 | 56 | 0.896 | 0.502(0.012) | 0.542(0.020) | 0.780 | 0.477(0.019) | 0.475(0.016) |
| 37 | 13 | 33 | 0.797 | 0.529(0.012) | 0.516(0.014) | 0.584 | 0.519(0.018) | 0.489(0.015) |
| 37 | 19 | 34 | 0.814 | 0.503(0.017) | 0.485(0.014) | 0.540 | 0.518(0.024) | 0.430(0.019) |
| 37 | 20 | 39 | 0.763 | 0.529(0.012) | 0.462(0.017) | 0.625 | 0.493(0.024) | 0.449(0.017) |
| 37 | 23 | 39 | 0.797 | 0.534(0.017) | 0.501(0.024) | 0.580 | 0.510(0.017) | 0.483(0.012) |
| 37 | 26 | 51 | 0.746 | 0.487(0.016) | 0.467(0.017) | 0.558 | 0.487(0.015) | 0.514(0.015) |
| 37 | 27 | 56 | 0.780 | 0.497(0.014) | 0.506(0.017) | 0.660 | 0.525(0.014) | 0.487(0.016) |
| 37 | 37 | 83 | 0.864 | 0.498(0.017) | 0.502(0.017) | 1.000 | 0.498(0.017) | 0.534(0.024) |

**Supplementary Table 1. Decoding performance across sessions using longitudinally track populations of individual neurons**. Within and across session decoding performance for each session pair chosen among 7 sessions spanning 24 days.

### Supplementary Videos

**Supplementary Video 1. Plug and play daily imaging with the head-mounted miniscope.**
See Supplementary Material to download.

**Supplementary Video 2. Cellular resolution calcium imaging in dorsal premotor cortex of behaving macaque.**
See Supplementary Material to download.

**Supplementary Video 3. Stable calcium imaging in head-unrestrained and behaving macaque.**
See Supplementary Material to download.

**Supplementary Video 4. Sedated blood flow imaging.**
See Supplementary Material to download.

**Supplementary Video 5. Calcium dynamics selective to contralateral arm reach direction.**
See Supplementary Material to download.

**Supplementary Video 6. Bilateral calcium dynamics selective to left and right arm reaches.**
See Supplementary Material to download.
